# Characterization of transcriptomic changes across *Coccidioides* morphologies using RiboMarker®-enhanced RNA sequencing

**DOI:** 10.1101/2025.02.11.634332

**Authors:** Jonathan M. Howard, Aidan C. Manning, Rachel C. Clark, Tahirah Williams, Clarissa J. Nobile, Sergei Kazakov, Sergio Barberan-Soler

**Affiliations:** RealSeq Biosciences, 2125 Delaware Ave Suite B, Santa Cruz, CA, 95060; Department of Molecular and Cell Biology, University of California, Merced, 5200 North Lake Road, Merced, CA, 95343; Quantitative and Systems Biology Graduate Program, University of California, Merced, 5200 North Lake Road, Merced, CA, 95343; Health Sciences Research Institute, University of California, Merced, 5200 North Lake Road, Merced, CA, 95343

## Abstract

*Coccidioides* is a dimorphic, pathogenic fungus responsible for transmission of the mammalian disease colloquially known as “Valley fever”. To better understand the molecular basis of *Coccidioides* pathogenesis, previous studies have characterized transcriptomes that define transitions between the saprobic and pathogenic life stages of the two species that cause Valley fever - *Coccidioides immitis and Coccidioides posadasii*. However, none of these studies have focused on small RNA profiles, which have been shown in several pathogenic fungi to play crucial roles in host-pathogen communication, affecting virulence and infectivity. In this study, we analyzed changes in small RNA expression across three major morphologies of *C. posadasii*: arthroconidia, mycelia, and spherules, from both intracellular and extracellular fractions. Utilizing RiboMarker® small RNA and RNA fragment library preparation, we show enhanced coverage across the transcriptome by increasing incorporation of normally incompatible RNAs into the sequencing pool. Using these data, we observed transcriptomic shifts during the transition of arthroconidia to either mycelia or spherules, marked largely by changes in both protein-coding, tRNA, and unannotated loci. As little is known regarding the mechanisms governing these life stage transitions, these data provide better insight into those small RNA- and fragment-producing genes and loci that may be required for progression between *Coccidioides* saprobic and parasitic life cycles. Additionally, analysis of fragmentation patterns across all morphologies suggests unique patterns of RNA fragmentation across a cohort of RNA species that correlate with a given ecotype. Finally, we noted evidence of RNA export to the extracellular space, particularly regarding snRNA and tRNA-derived fragments as well as mRNA-derived transcripts, during the transition to either mycelia or spherules, which may play roles in cell-cell, and/or host-pathogen communication. Going forward, this newly established intra- and extracellular *Coccidioides* sRNA atlas will provide a foundation for potential biomarker discovery and contribute to our understanding of the molecular basis for virulence in Valley fever.

## INTRODUCTION

*Coccidioides* (which includes *C. immitis and C. posadasii)* is a genus of dimorphic fungi endemic to the dry and desert regions of the Southwestern United States and Mexico. Together, they are responsible for the disease colloquially known as “Valley fever” (or coccidioidomycosis) (1). Latest data indicate *Coccidioides* is responsible for nearly 20,000 clinically significant cases of Valley fever per year in the U.S. (2), a number that is likely underestimated due to misdiagnosis, underreporting, and lack of medical intervention (3). Furthermore, estimates predict an increasing prevalence of Valley fever throughout much of the Western U.S. due to the more general, emerging threats of antifungal-resistant pathogenic fungi (4), a growing immunocompromised population (5, 6), as well as environmental factors, such as climate change (7). Consequently, the economic ramifications of coccidioidomycosis are expected to increase from $3.9 billion in 2015 to over $18 billion by the end of the 21st century (8). As of 2023, the World Health Organization’s (WHO) inaugural list of “Fungal Priority Pathogens” included *Coccidioides* as a microbe of growing concern, requiring increased attention concerning basic research and the development of clinical tools (both diagnostic and pharmacological) (9).

To tackle these questions, researchers are providing insights into the modulation of transcriptional programs that support the saprobic-to-parasitic phase transition of *Coccidioides* using next-generation sequencing (NGS) approaches (10). RNA sequencing experiments comparing mutant strains and wild-type strains of *C. posadasii* suggest transition to spherule (parasitic) morphology is accompanied by increased expression of transcripts associated with nutrient assimilation, most likely as a response to host milieu (11). Furthermore, by assessing nascent global transcriptomes to examine differentially expressed genes in the saprobic and parasitic life cycles of *C. immitis* (12), Duttke et al. detected expression changes in genes related to stress response, cell wall remodeling, polar growth, transcription factors, known virulence factors, and other genes of interest. Many of these gene expression changes were associated with known *cis*-regulatory sequences utilized by the conserved transcription factor Ryp1, a key player in developmental transition conserved across the fungal kingdom (13). Overall, these data and others provide a foundation to understanding the causative transcriptional programs of saprobic-to-parasitic transition in *Coccidioides*, as well as a list of putative targets for diagnostic and therapeutic purposes to ameliorate the impending spread of Valley fever.

While these data have been informative to our work herein, they overlook or select against sequencing of the small noncoding (sRNAs) and fragmented pools of RNA. sRNAs are defined by species of RNA less than 200 nucleotides in length, harboring distinct subclasses (e.g. tRNAs, miRNAs, snRNAs, etc.) with a variety of known biological functions (14). By definition, sRNAs also include fragments of both large and small RNAs, often generated by exo- and endonucleolytic cleavage, and themselves may represent types of sRNAs (e.g. tsRNAs or tRFs, rRFs) (15, 16) or products of regulated processing and/or degradation (17). In the context of fungal pathogens, sRNAs have been shown to be key mediators of pathogenicity. For example, the fungal-like pathogen *Phytophthora sojae* (the cause of stem and root rot in soybeans) generates small RNAs to mediate transgenerational gene silencing of the avirulence gene *Avr3a* to increase pathogenicity and evade host immune surveillance in progeny (18). Additionally, the microRNA-like RNA (milRNA) VdmilR1 in *Verticilium dahlae*, a soil-borne fungal pathogen responsible for wilt disease, targets the putative protein-coding gene VdHy, an essential gene regulating pathogenicity through chromatin silencing (19). Therefore, sRNAs can offer an additional source of information to glean insight into dynamic biological processes and gene regulation as they modulate over pathogen life cycles and during infection.

Beyond the intracellular space, various sRNAs have been found to populate extracellular vesicles (EVs) exported by fungal pathogens. Extracellular vesicle (EV) biogenesis and trafficking play critical roles in delivering concentrated payloads of fungal macromolecules to host effector cells and tissues (20). These mechanisms are largely thought to rely on EV-transported miRNAs/milRNAs and siRNAs from pathogen to host, and vice versa, to modulate host gene expression and facilitate infection (in the case of pathogen to host) or mitigate virulence (in the case of host to pathogen) (21–23). For example, the fungal pathogen *Botrytis cinerea* can export small RNAs to enter plant cells and hijack silencing factors to dampen immunity-related genes (24). Similar pathways have been discovered between the interaction of the pathogenic fungus *Beauveria bassiana* and its host, the *Culicidae* (mosquito), through exporting the small RNA bba-milR1 to inhibit immunity factor Spz4 and facilitate infection (25). With these data, the transport and release of miRNAs/milRNAs into nearby host tissues have been shown to contribute, in part, to the pathophysiological regulation of fungal infections, most notably being cross-kingdom RNA interference (26, 27). However, many subtypes of sRNAs (e.g. snoRNAs, tRNAs/tRFs, snRNAs) are found to populate fungal EVs, many with canonical functions outside of the direct suppression of mRNA translation (28). By identifying what RNAs are packaged in these EVs of *Coccidioides* is not only a first step towards uncovering their putative roles in host-pathogen interactions, but also an additional layer informing the discovery of RNA biomarkers, (especially those found outside the cell), prognostic of Valley fever infection.

To better characterize the small RNA profile of *Coccidioides,* we performed small RNA sequencing using an enhanced RealSeq-RiboMarker® protocol to generate the most complete “small RNA atlas” of *C. posadasii.* We profiled across a 96-hr growth time course of cultures for the saprobic mycelia and arthroconidia as well as from parasitic spherules. Additionally, we implemented the RealSeq RiboMarker® method to reveal additional layers of full-length and fragment sRNAs often unincorporated into sequencing libraries. Furthermore, we interrogated cell-free and exosomal RNA from conditioned media for these morphologies to determine what, if any, small RNAs may be exported from the fungus into the extracellular space. Our data reveal that morphological changes are not only driven by protein-coding genes, but also by annotated sRNA types, including tRNA fragments (tRFs/tDRs), as well as unannotated sRNA loci. Additionally, we identified the export of key extracellular vesicle-associated RNAs which may define and shape cell:cell, and pathogen:host interactions. Ultimately, this work strives to further characterize the distinct RNA profiles associated with these important life stage morphologies of *Coccidioides* and provide a foundational resource of potential RNA biomarkers to target in *ex vivo* samples of infected hosts.

## MATERIAL AND METHODS

### Culture Conditions and Harvesting

*C. posadasii* (strain Δ*cts2*/Δ*ard1*/Δ*cts3*, NR-166; BEI Resources) arthroconidia were harvested from 6-week-old plates as previously described (29). The spores were inoculated at 1 × 10^6^ arthroconidia into a 250 mL baffled Erlenmeyer flask containing 100 mL of 2xGYE, shaking at 150 rpm at 30 °C for 48 hours. Conditioned media was harvested by collecting media with cells into centrifuge tubes, which were centrifuged at 12,000 × *g* for 8 min to pellet the cells. The supernatant was pooled and filtered with a 0.22-µm syringe filter before being stored at −80°C.

To generate samples for the time course, in which arthroconidia transition to mycelia, 1 × 10^6^ of harvested arthroconidia were incubated in 250 mL flat-bottom Erlenmeyer flasks (Corning, Corning, NY, USA) in 100 mL of 2x GYE media, shaking at 150 rpm at 30 °C. After 48 hours, the conditioned media with cells (predominantly in the arthroconidia morphology) were centrifuged at 12,000 × *g* for 8 min to pellet the cells. Half of the filtered fungal cells were kept for RNA isolation, while the condition media was spun down, and supernatant was removed for extracellular RNA isolation. The remaining cells were reinoculated into fresh 2X GYE for an additional 24 hours, shaking at 150 rpm at 30 °C. At this 72-hour time point, the fungal cells (containing a mixture of both arthroconidia and mycelia morphologies) were separated from the growth media by filtering with a sterile 40 µm nylon mesh cell strainer. Half of the filtered fungal cells were kept for RNA isolation, while the conditioned media was filtered with a 0.22-µm syringe filter before being stored at −80°C for extracellular RNA isolation. The remaining half of the filtered fungal cells were reintroduced again into fresh 2X GYE for 24 hours, shaking at 150 rpm at 30 °C. This sample processing was performed again for the cell samples at the 96 hours time point (predominantly in the mycelia morphology) for a total of 3 replicates of time course fungal cell samples and filtered conditioned media.

*C. posadasii* spherules were propagated and harvested from 6-week-old plates as previously described (29). For spherule samples, 50 mL of Converse medium (30) was inoculated to a final concentration of 10^6^ arthroconidia/mL. Cultures were then incubated at 39°C in 10% CO_2_, shaking at 150 rpm for ∼ 5 days. Spherules were harvested by filtering the culture through a nested filter into a 50 mL conical tube and were then centrifuged at 9000 x *g* for 8 min to pellet the cells. Cell pellets were washed twice with 1x PBS and then snap frozen for storage at −80°C until use.

### Fungal and extracellular RNA extraction and purification

*C. posadasii* mycelia, arthroconidia, and spherules samples were stored in RNA Shield (Zymo Research) at −80°C until processing using the Quick-RNA™ Fungal/Bacterial Microprep kit (Zymo Research) according to the manufacturer’s instructions. Samples were added to a pre-chilled 2 mL ZR Bashing Bead lysis tube with 0.5 mm beads (Zymo Research), and tubes were arranged in a FastPrep-24™ grinder and lysis system (MP Biomedicals) and disrupted 5x for 60 s at 6.5 m/s, with intervening incubations on ice for 5 min. DNase I treatment on-column was also implemented. Cell-free and exosome RNAs from *C. posadasii* mycelia and arthroconidia samples were isolated from 40 mL of conditioned culture media using the Plasma/Serum Exosome and Free-Circulating RNA Isolation Kit (Norgen Biotek) according to the manufacturer’s instructions. Eluted total and extracellular RNAs purified from mycelia, arthroconidia, and spherules samples (1 replicate per stage), as well as conditioned media, were quantified using a Qubit 3.0 fluorometer (Invitrogen).

### RNA pretreatment with RiboMarker®^®^

For treatment of RNA with the RiboMarker® platform, 10 µL of RNA sample was incubated in a 15 μL reaction mixture containing 1.5 μL RiboMarker®^®^ Buffer 1, 1.5 μL of RiboMarker® Enzyme 1, and 2 μL of RNase-Free H_2_O at 37°C for 10 min, followed by 80°C for 2 min. Next, 2 μL of RiboMarker® Buffer 2 and 2 μL of RiboMarker® Enzyme 2 was added to the mixture, which was then incubated at 37°C for 60 min. Finally, 1 μL of RiboMarker® Buffer 3 was added to the reaction and incubated at 37°C for 60 min. The pretreated RNA was isolated using an RNA Clean and Concentrator-5 kit (Zymo Research) for downstream use in library preparation.

### Library Preparation and Sequencing

For small RNA-seq, libraries were prepared from total RNA using the RealSeq^®^-Biofluids library preparation kit (RealSeq Biosciences) according to the manufacturer’s instructions. Libraries for each ecotype, as well as cell-free and exosome samples, were generated separately and then pooled for the analysis. In some cases, triplicate libraries for each sample type and condition were not met, and thus, only duplicates were used in downstream analyses to avoid batch effects in the data. Finally, prepared libraries were pooled and subjected to single-end 75 cycles of sequencing using the F3 50 cycle sequencing kit (Note: reagents for 50 additional cycles above what is represented were included to account for index sequencing needs) on the G4 system according to the manufacturer’s instructions (Singular Genomics).

### Data Analysis

Sequencing reads were trimmed of adapter sequence using cutadapt version 4.6 [--nextseq-trim=15 -u 1 -a TGGAATTCTCGGGTGCCAAGG -m 15] (31). Reads were then aligned to the *C. posadasii* strain Silveira (GCA_000170175.2) with the addition of the high-confidence set of tRNA isodecoders containing -CCA tails identified using tRNAScan-SE (32). ShortStack v4.1.0 (33) with the parameters [-pad 3 –mincov 0.5rpm] was used to identify potential novel sRNA transcripts. Read counts were normalized using DESeq2 version 1.42.0.

### RNA Fragmentation Analysis

Read pileups across the body of non-coding RNA (ncRNA) transcripts, from 5′ to 3′, were extracted and converted into a linear vector representing read coverage across each molecule. Coverage was normalized to the total number of reads mapping to each transcript, and a one-way functional ANOVA (34) was applied to assess significant differences in coverage patterns across morphologies, using a threshold of p < 0.05. To further explore these patterns, we employed a probabilistic functional data clustering approach, specifically fuzzy K-means, to classify transcripts based on their normalized read coverage profiles. The model was trained with three clusters corresponding to the distinct Coccidioides life morphologies: Arthroconidia, Mycelia, and Spherule. Each sample was subsequently assigned probabilities reflecting the strength of association between its coverage pattern and the representative functional cluster for each morphology. For each transcript a “cluster_score” was calculated that reflected the strength of the fuzzy k-means approach for identifying morphology-specific fragmentation patterns with a score of 1 indicating perfect morphology-driven patterns, and less than 1 indicating samples from different morphological stages had similar fragmentation patterns.

## RESULTS

### RiboMarker®-Enhanced RNA sequencing of *Coccidioides* morphologies

Most conventional small RNA (sRNA)-Seq library preparation kits are limited to the capture of RNA species with 5’-phosphate, 3’-hydroxyl, and minimal (if any) internal base modifications. Recent data identifies only a small portion (∼10%) of any small RNA transcriptome may be endogenously compatible (35), suggesting many of these RNA species are not adequately captured using commercially available library preparation kits. To address these limitations, we investigated potential improvements to pretreatments collectively referred to as “end-healing”, or the chemical/enzymatic conversion of incompatible RNA ends to those required for adapter ligation during library preparation. T4 Polynucleotide Kinase (T4PNK) is a commonly used enzyme to achieve compatible ends due to its dual 5’ kinase and 3’ phosphatase activities (36, 37), however, the optimum conditions for PNK reactions have not been determined when detecting naturally occurring RNA fragments.

To interrogate the capacity of T4PNK to increase RNA inclusion in sequencing libraries, we tested different T4PNK pretreatments under conditions previously described (37), to determine their respective effects on the sequencing output. We assessed four different buffers: the RiboMarker® Buffer (RealSeq Biosciences), imidazole, TRIS (pH 6.0), and the standard (NEB formulated) T4PNK Buffer supplied with the enzyme, and performed experiments in the presence and absence of 10 µM ATP (or 1 mM ATP for standard conditions as suggested by the manufacturer), as sustained T4PNK activity may require the presence of this substrate (37, 38). From the resulting sequencing data, we determined the number of unique RNA transcripts identified in each sample as a measure of RNA diversity by subsampling 5 million reads per sample and marking a transcript as “detected” if it had at least 10 mapped reads. These results (Supplemental Figure 1) revealed appreciable changes to the detected RNA classes with respect to the differential PNK reaction conditions (“None” vs. all treatments). However, T4PNK treatment in the presence of RiboMarker® Buffer resulted in two of the most notable differences compared to other conditions. We observed a distinguishable increase in protein-coding gene-derived RNAs using RiboMarker® conditions (Supplemental Figure 1, “RiboMarker Buffer”), as well as a slight, but detectable increase in all other classes, apart from rRNAs (ribosomal) and scaRNAs (small Cajal-body-specific RNAs). To further validate the ability of RiboMarker® Buffer to increase RNA inclusivity, we sequenced a pool of synthetic spike-ins (see Supplemental File 1) alongside these natural RNAs, which contained all potential end type combinations (Type 1 = 5’ Phosphate, 3’ Hydroxyl; Type 2 = 5’ Hydroxyl, 3’ Hydroxyl; Type 3 = 5’ Hydroxyl, 3’ Phosphate; Type 4 = 5’ Phosphate, 3’ Phosphate). Libraries were prepared using the RealSeq-Biofluids kit, both without RNA pretreatment (“BioFluids”) and with T4PNK treatment in RiboMarker® Buffer (“RiboMarker®”), and a library using an alternative competitor small RNA library preparation kit with standard T4PNK treatment in the presence of ATP (“Phospho-Seq”). These results (Figure 1A) confirmed that T4PNK treatment in RiboMarker® Buffer resulted in the inclusion of a more diverse set of RNA molecules containing both compatible and previously incompatible RNA ends into the sequencing library and yielded a more evenly balanced read distribution for all four RNA end types vs. the alternative kit with standard T4PNK treatment. Therefore, these data revealed that T4PNK RNA pretreatments, in the presence of RiboMarker® Buffer, facilitated the most unbiased, and therefore diverse, inclusion of RNA molecules into resulting libraries.

**Figure 1.**
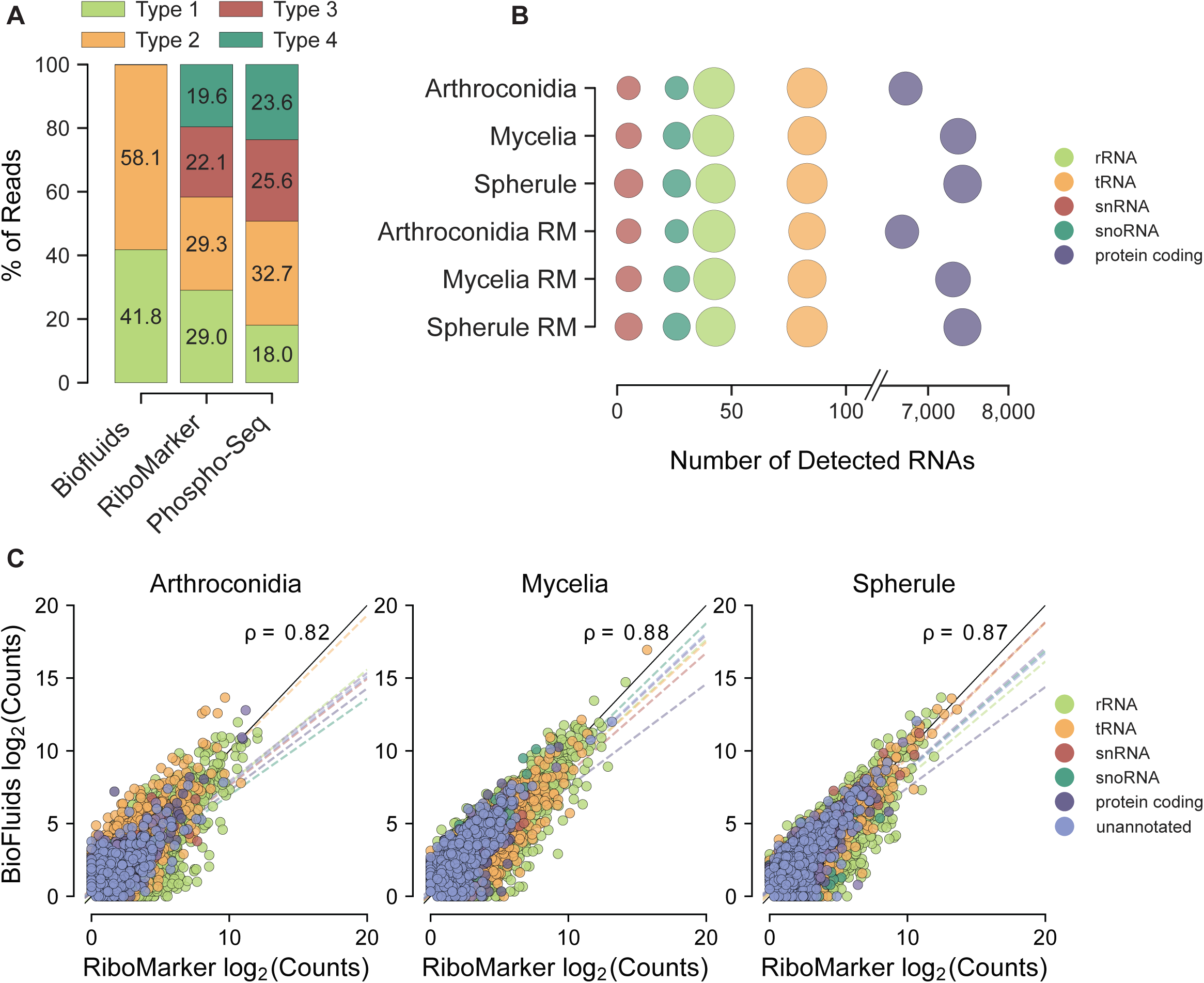
RNA sequencing across 3 *Coccidioides* morphologies of *C. posadasii.* **(A)** Bar plot showing percentages of spike-ins representing various RNA end types detected in libraries prepared with different methods: RealSeq^®^-Biofluids Library Prep without RNA pretreatment (“Biofluids”), RealSeq^®^-Biofluids Library Prep with T4PNK RNA pretreatment in RiboMarker® Buffer (“RiboMarker®”), and NEBNext^®^ Small RNA Library Prep with standard PNK pretreatment (“Phospho-Seq”). For Biofluids, Type 3 and 4 reads are also below 1% and are thus not visible. Types are defined as follows: Type 1 = 5’ Phosphate, 3’ Hydroxyl; Type 2 = 5’ Hydroxyl, 3’ Hydroxyl; Type 3 = 5’ Hydroxyl, 3’ Phosphate; Type 4 = 5’ Phosphate, 3’ Phosphate. (**B**) Dot plot representing the number of detected transcript annotations mapped to C. posadasii in Biofluids (top) and RiboMarker® (bottom) libraries, with dot size proportional to the abundance of each RNA class. (**C**) Scatter plot of the abundance of non-redundant transcripts between Biofluids and RiboMarker® for an arthroconidia (left; Pearson=0.82), mycelia (middle; Pearson=0.88), and spherules (right; Pearson=0.87).

To expand upon the use of RiboMarker® into a new sequencing space, we chose to characterize the dynamic changes in small RNA expression across the multiple morphologies of *Coccidioides*, an epidemiologically important, yet understudied, fungal pathogen (39). We utilized *C. posadasii*, Δ*cts2*/Δ*ard1*/Δ*cts3*, NR-166 as our model strain (40), and harvested arthroconidia, mycelia, and spherules samples, representing several key morphologies for this important fungal pathogen. Arthroconidia were inoculated into growth media and grown for four days, with timepoints harvested at 48 hrs (arthroconidia) and 96 hrs (mycelia), representing the transition of *Coccidioides* arthroconidia to mycelia. Additionally, to generate spherules, the same arthroconidia were harvested after growing for 5 days in Converse media (30). For assessing the presence of potential extracellular *Coccidioides* RNAs, conditioned growth media from all morphologies was also collected and used for cell-free RNA (cfRNA) and exosomal RNA isolation. Additionally, each sample was subjected to two different RNA pretreatment conditions: one utilizing no RNA pretreatment and one implementing the optimized T4PNK pretreatment in RiboMarker® Buffer (Figure 1A). Together, these two datasets enabled us to comprehensively evaluate the small RNA transcriptome of each morphology tested as well as ascertain what differences, if any, in RNA expression and fragmentation occurred across the saprobic and parasitic life cycles of Coccidioides.

The resulting sequencing libraries were processed (see Methods) and the differences across the morphologies of *C. posadasii* were explored using both Biofluids and RiboMarker® library preparations. An initial assessment of the read lengths suggested that RiboMarker® reduces biases to specific RNA types (Supplemental Figure 2). For example, the arthroconidia samples (Supplemental Figure 2A; “Arthroconidia”) exhibited large peaks at ∼ 28 nt, often indicative of the inclusion of specific RNAs, while a more even distribution from ∼15 to 35 nts was observed with RiboMarker® pretreatment. Read length patterns for intracellular spherules (Supplemental Figure 2A; “Spherule”) looks similar between “BioFluids” and “RiboMarker”, with the exception of a more pronounced peak at ∼35 nts, possibly indicative of tRNA halves or fragments (41). Looking at the pool of distinct RNA molecules captured from each morphology in our panel using both Biofluids and RiboMarker® T4PNK pretreatment (Figure 1B and C), we found a strong correlation between these library preparations (Figure 1C). However, these data also point to the enrichment of key sRNA sub-populations in the RiboMarker® samples that may otherwise not be detected using no treatment approaches (Figure 1C; hatch marks). Indeed, we noted large increases in reads derived from ribosomal RNA (rRNA) across the arthroconidia and mycelia samples, as well as tRNA-derived reads in the spherules samples (Supplemental Figure 3A), suggesting that RiboMarker® treatment led to a shift in the incorporation of previously incompatible RNA ends that would otherwise be biased against (42). An assessment of the overlapping molecules captured using either Biofluids or RiboMarker®, (Supplemental Figure 4) revealed ∼50-60% RNAs were incorporated using either method, while the remaining ∼40-50% of reads showed specific inclusion in either Biofluids (∼13-16%) or RiboMarker® (∼24-33% of all unique reads) preparations. By and large, we observe minimal differences in the detected expression across most annotated sRNA gene types (Figure 1B), regardless of RNA pretreatment. However, we did observe differences in the number of reads mapping to *Coccidioides* protein-coding genes, especially in the spherules samples, suggesting that these sRNAs may be specifically upregulated, or are a byproduct of degradation/processing of larger mRNA species turned on during transition to this morphology (10). Using a principal-component analysis (PCA; Supplemental Figure 5), we observe that regardless of the library preparation method, clear separation of each time point suggested distinct populations of small RNAs/RNA fragments were captured in the different *Coccidioides* morphologies. Taken together, these data suggest there are sub-populations of RNA that are more efficiently captured using RiboMarker®, resulting in a more diverse, and unbiased, sequencing library. Furthermore, these results highlight that to completely capture the diversity of small RNA molecules found in each biological sample, multiple approaches may be needed, and using either approach, we note distinct small RNA profiles across *Coccidioides* life stages.

### Characterization of *Coccidioides* sRNA profiles from distinct morphologies reveals ecotype-specific differences in expression

While the transition of arthroconidia to mycelia or spherules is largely dependent on environmental factors, such as changes in CO_2_ levels (43), recent literature has begun to tease apart the transcriptional reprogramming required to support the external cues that trigger transitions between *Coccidioides* morphologies (10). We sought to further our understanding of the transcriptional changes driving *C. posadasii* morphological transitions using RiboMarker® small RNA sequencing.

We observed differences in detected proportions of sRNA transcript types across sample types in the early time points (Figure 2A). For example, we see small, but noticeable increases in tRNA-derived and protein-coding-derived sRNA detection in transitions between arthroconidia to mycelia. This may be indicative of similar small RNA expression levels among saprobic stage samples, but also reflecting differences with respect to morphology, especially with increases in transcription and translation. We also observed these same expression changes magnified with respect to spherules, suggesting a similar increase in gene expression which may be representative of transcriptomic and translational changes during the transition to the parasitic life cycle (10). A deeper look at ncRNAs detected in our study suggest that as compared to protein-coding derived RNAs, of which only 50% appear to change in expression across all life cycle morphologies (Figure 2B, “protein coding”), we find that nearly 80% or more of annotated ncRNAs subtypes detected in our data show perturbations. This suggests large transcriptomic profile modulations, coding and non-coding, are a defining characteristic of morphologies in *Coccidioides*.

**Figure 2.**
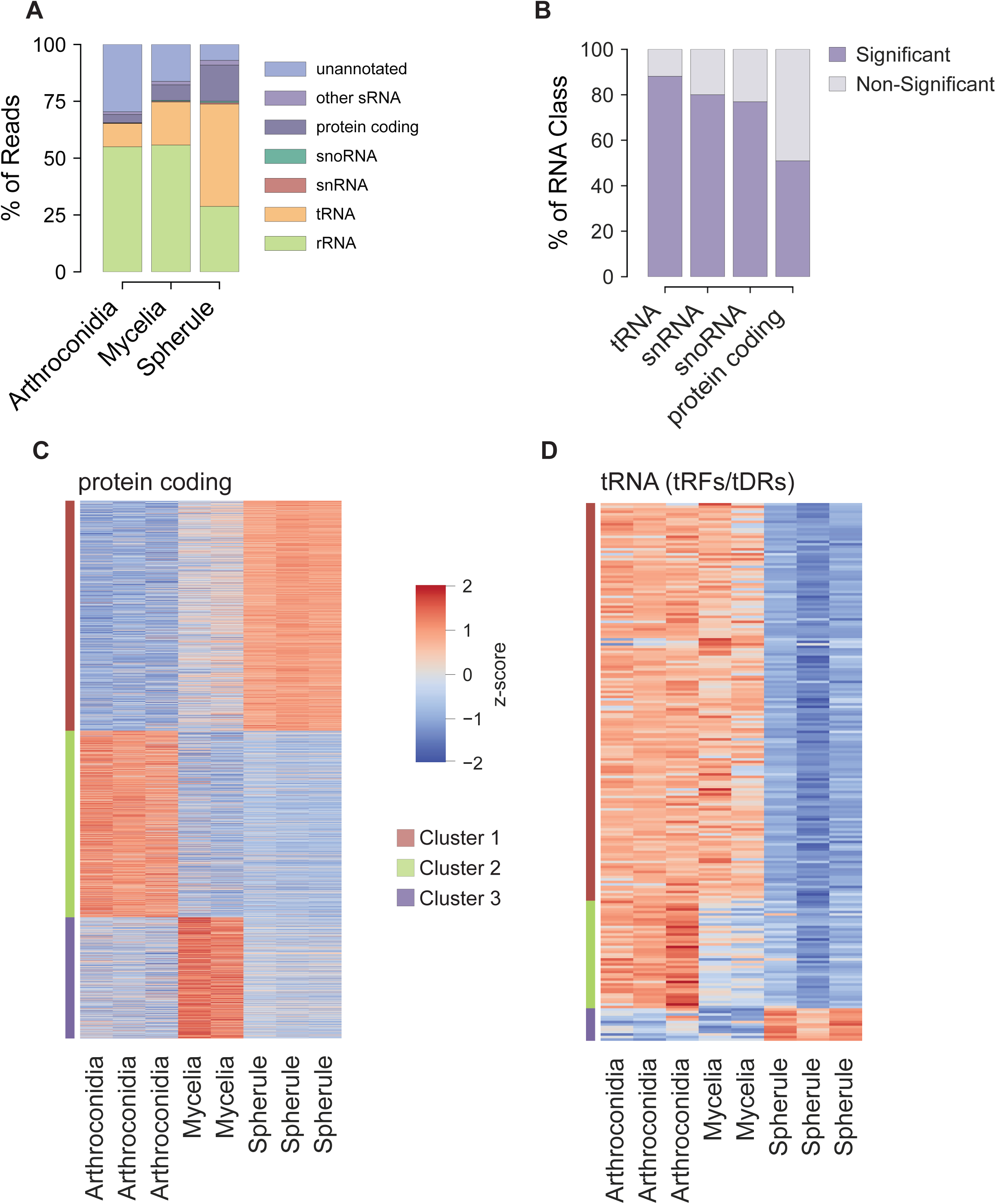
Differential expression analysis of annotated RNAs across *C. posadasii* morphologies. (A) Bar plot showing the percentage of mapped RiboMarker® small RNA reads categorized by RNA type annotations in *C. posadasii*. (B) Bar plot showing the percentage of transcripts from different RNA classes found significantly enriched (padj<0.01) in a *Coccidioides* morphology. **(C, D)** Hierarchical clustered heat maps of protein-coding RNA abundance (**B**; 3,474 significantly different [padj < 0.01]) and tRNA fragments (**C**; 215 significantly different across time points [padj < 0.01]) in RiboMarker® intracellular small RNA sequencing datasets. The color gradient represents Z-scores, with clusters indicated by colored bars on the left.

An assessment of the abundance of small RNA molecules derived from mRNAs across our samples identified 3,641 that significantly fluctuated during the time course (Supplemental Table 2). A heatmap of these is presented in Figure 2C (data in Supplemental File 2) where we observed limited variation in transcript abundance among replicates for each morphology. Notable patterns of expression were made evident through hierarchical clustering of these data where we were able to define four clusters specific to the arthroconidia (Figure 2C, cluster 2), mycelia (Figure 2B, cluster 3), and spherules (Figure 2B, cluster 1) ecotypes. From each of these clusters, we isolated representative examples which highlight the differences in mRNA-derived small RNAs abundance across time points (Supplemental Figure 6). For example, we saw a relatively high abundance of small RNAs generated from the gene *INDA1* specific to mature mycelia (Supplemental Figure 6A). This gene encodes an amino acid permease, responsible for, among other processes, the uptake of amino acids from the environment (44, 45). Regarding spherule enriched expression, we saw increased capture of reads derived from *HSK1* (Supplemental Figure 6C), which encodes a serine-threonine kinase, whose homologue has been shown to regulate DNA replication initiation in fission yeast (46), a process known to be highly upregulated in the spherule morphology (47). Finally, we identified two transcripts differentially expressed along the arthroconidia to mycelia transition, one that encodes 4-hydroxyphenylpyruvate dioxygenase (HPPD; Supplemental Figure 6B), a protein responsible for the metabolism of tyrosine and phenylalanine as a carbon source (48), and the other that encodes Sec24, a Coat Protein Complex II (COPII)-associated protein (Supplemental Figure 6D), responsible for the intracellular transport of ER-derived proteins throughout the fungal cell (49).

Another subtype of RNA that revealed substantial changes in abundance were fragments derived from transfer RNAs (or tRFs/tDRs). A heatmap depicting the expression levels of differentially expressed tRNA-derived RNAs is presented in Figure 2D (data in Supplemental File 3). Along with the high degree of reproducibility among morphological replicates, we also noted that the abundance of tRNA fragments was generally high in the arthroconidia and mycelia relative to the spherules (Figure 2D, clusters 1 and 2). This may be reflective of the observed differences in protein synthesis among morphologies (50). By and large, spherules showed low levels of tRNA fragment abundance (Figure 2D, cluster 1 and 2, spherules samples) compared to arthroconidia and mycelia, suggesting a higher requirement for full-length tRNAs to regulate protein synthesis (51). Interestingly, spherules contained a small cluster of tRNA fragments that were specifically abundant (Figure 2D, cluster 3), including an abundance of Glycine and Tyrosine tRNA 3’ halves, Histidine 5’ halves, and Serine 3’-derived fragments (Supplemental Figure 7). While previous literature has shown tRNA fragmentation to globally regulate protein translation, mRNA stability, and RNA binding protein activity (52), their functions in fungi have remained elusive.

### Transcriptomic analysis reveals distinct populations of unannotated small RNA producing loci

Beyond the confines of annotated RNAs derived from the *Coccidioides* genome, our transcriptional data suggests a small but significant population of RNA is derived from unannotated (or intergenic) regions (Figure 2A). To identify potential novel RNA producing loci, we used reference-aligned, RiboMarker® RNA sequencing reads as input into ShortStack (*Methods*; 33), a bioinformatics tool that performs comprehensive *de novo* annotation and quantification of inferred small RNA genes (regardless of annotation) and fragment-generating loci. These ShortStack-inferred RNA loci were then mapped against the *C. posadasii* genome to assess where they land relative to its annotated and unannotated regions. Results from our ShortStack analysis revealed that a subset of small RNA read clusters were derived from rRNAs and tRNAs (Figure 3A), while most inferred clusters were found in protein-coding genes (89.2%), with unannotated regions comprising the second highest population (6.2%; Figure 3A). Granted, while clusters mapping to rRNA and tRNA genes are high in average read count (Figure 3B), which is to be expected, unannotated make up a smaller but more diverse cohort of detected RNAs (Figure 3D). While this RNA sub-population constitutes what is often described as the “dark matter” of the genome, this phenomenon is found across all branches of life (53, 54), including fungi (55). Identification of these seemingly novel RNAs is further complicated by the current state of genomes from *Coccidioides* and other fungi, as they remain poorly characterized, leading to less properly annotated coding and non-coding genes. Therefore, to further characterize these unannotated-derived RNA species, we took a data-driven, unbiased approach to identify loci of unannotated RNA expression across the morphologies of *Coccidioides* central to this study.

**Figure 3.**
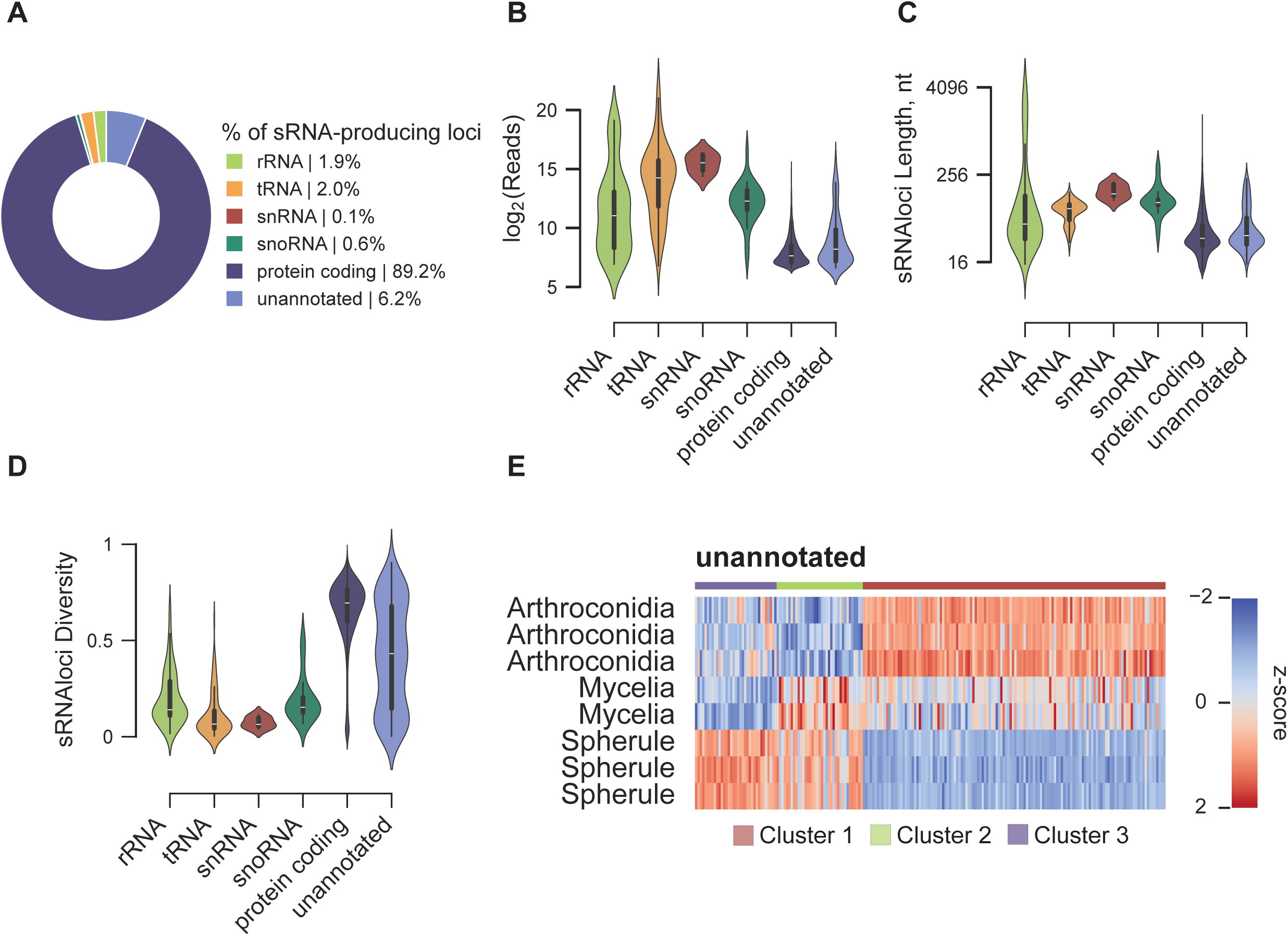
Characterization and differential expression of unannotated RNA loci across *C. posadasii* morphologies. **(A)** Pie chart illustrating the distribution of ShortStack clusters within each annotated RNA type or if it fell within an unannotated region. (**B/C/D**) Violin plots showing the (B) log₂ average read counts of mapped ShortStack loci by RNA type; (C) length of ShortStack loci identified among the different RNA classes; (D) diversity of reads within each ShortStack loci. (**I**) Clustered heatmap of unannotated ShortStack loci showing significant differential expression across time points (padj. < 0.01) in both intracellular small RNA samples. Cluster groups are indicated by colored bars at the top.

To further characterize the unannotated loci from our RiboMarker® *Coccidioides* small RNA libraries, we defined them using criteria described in Dorga et al. (56) (*Methods* & Figure 3B–D) using the following metrics: locus read abundance (Figure 3B), locus length (Figure 3C), and locus read diversity (Figure 3D). We observed significant differences (Figures 3B and 3C; p<0.0001; *Methods*) in locus abundance and length when comparing annotated non-coding RNAs (ribosomal RNAs [rRNAs], transfer RNAs [tRNAs] small nuclear RNAs [snRNAs], and small nucleolar RNA [snoRNAs]) to both protein-coding-derived and unannotated RNA loci. This is reflective of both the bias of the library preparation for smaller RNAs and expression differences, especially between protein-coding and noncoding RNAs highlighted here. Interestingly unannotated clusters, along with protein-coding loci, show higher complexity (Figure 3D; pval. 1×10^−253^), compared to annotated ncRNA clusters. These data suggest that ShortStack small RNA loci that fall within annotated regions (such as protein-coding genes) generally show less diverse coverage than unannotated loci, which may be representative of a lower-complexity region or high-specificity in fragments that are generated from ncRNAs in general. Additionally, unannotated genes may, in part, reflect both coding and non-coding genes that are yet to be annotated, possibly explaining why unannotated regions are “diverse” in read diversity.

Similarly for annotated transcripts, we identified RNAs produced from these novel loci that significantly fluctuated during the life cycle of *C. posadasii* and can be found in Figure 3E (data in Supplemental File 4). As with annotated regions, hierarchical clustering revealed subsets of these transcripts that exhibit morphology specific expression (Figure 3E; Clusters 1-3; logFC2 values for this heatmap can be found in Supplemental Table 3). Focusing on the spherule morphology, we highlighted two examples (of many), which showed high levels of enrichment in this parasitic growth phase (Supplemental Figure 8). For example, downstream of CPSG_01053 (predicted to encode a CORD and CS-domain containing protein) (57), we identified sRNAlocus_4461 a novel small RNA-producing loci (Supplemental Figure 7A) whose expression is derived from the opposite strand of the neighboring gene. RNACentral (58) and nucleotide BLAST (59) sequence analyses of this (and fragments of this) locus returned no significant similarities to any known annotated RNA, fungal or otherwise, but rather a collection of non-coding and hypothetical RNAs (data not shown). An RNAfold analysis (60), however, revealed a potential secondary structure with a minimum free energy of −26.20 kcal/mol (Supplemental Figure 8B) representing a sequence from the most highly expressed regions of this cluster (Supplemental Figure 8A; highlighted red box). For context, analysis of 20 randomly generated RNA sequences with identical %GC content yielded an average of −15.97 kcal/mol with a standard deviation of ± 3.80 (data not shown). Another example can be found downstream of two protein-coding genes (on opposite strands), CPSG_03826 (encoding a 3-hydroxyisobutyryl-CoA hydrolase), an enzyme responsible for amino acid and fatty acid catabolism (61) and CPSG_03827 (encoding a predicted DUF1264-domain containing protein), a protein domain found in enzymes involved in polycyclic aromatic hydrocarbon biodegradation (59) and histone acetylation (62). Additionally, we identified sRNAlocus_4580, which potentially encodes three separate small RNA-producing loci of varying lengths (Supplemental Figure 8C) expressed in both the 96 hr mycelia time point and in spherules. Again, RNAcentral (58) and nucleotide BLAST (59) sequence analyses of this (and apparent fragments of this) cluster returned no significant similarities to any known annotated RNAs. An RNAfold analysis (60) of the two small RNA producing sections of sRNAlocus_4580 (Supplemental Figure 8C; highlighted red boxes) as well as the entirety of sRNAlocus_4580 revealed three intriguing potential secondary structures with a minimum free energy of −25.00, −23.10, and −53.40 kcal/mol, for Supplemental Figures 8D, E, and F, respectively.

While these data should not be taken as definitive proof of unannotated RNAs functionality, their putative structure and morphology-specific expression may suggest regulation of their expression and downstream utilization. To further validate the presence and expression patterns of unannotated RNAs across morphologies, we performed additional total RNA sequencing, (which includes RNAs from both small and long [> 200 nts] species), to detect example RNAs presented herein (or their parental RNAs) as well as assess the correlation of unannotated small RNA expression to total RNA expression. In general, when comparing RiboMarker-enhanced libraries to total RNA sequencing libraries derived from the same sample, consistently high correlation was found (Supplemental Figure 9) regardless of same-sample comparison. Additionally, we were able to validate that all example RNA loci shown herein were recapitulated in total RNA data, at least in the context of expression from said loci (data not shown). Taken together, these, and other, unannotated small RNA loci may contain functional elements, either consisting of yet-to-be canonical non-coding RNAs or potential microRNA-like RNAs (25, 63). Further work will be required to validate and ascertain the functions of these putative small RNAs.

### Differential fragmentation patterns of annotated and unannotated noncoding RNAs correlate with *Coccidioides* morphology

Previous literature has utilized RNA expression/abundance (64) and modification patterns (65, 66) as a means of categorizing biological sources, such as tissue identification or diseased vs. healthy samples (67), among others. While the molecular function of RNA fragments is poorly understood, data suggests these may, in part, be representative of protein-bound protected regions of RNA species, evidence of secondary structure, or regulated effectors of gene expression (68–70). Together, these data suggest that perturbations to cellular homeostasis or development-related gene expression, may, in turn, result in unique RNA fragment profiles reflective of these changes. For example, as shown in Figure 4A, assuming the same transcript in two different fungal morphologies, interactions with RNA binding proteins, modifications states of the RNA, flexible secondary structures, or regulated translation of mRNAs, one may hypothesize that these *cis-* and *trans-*acting effectors could lead to different processing or degradation of a given RNA. Therefore, we questioned whether RNA fragmentation patterns could be used as a classification system to identify *Coccidioides* morphologies.

**Figure 4.**
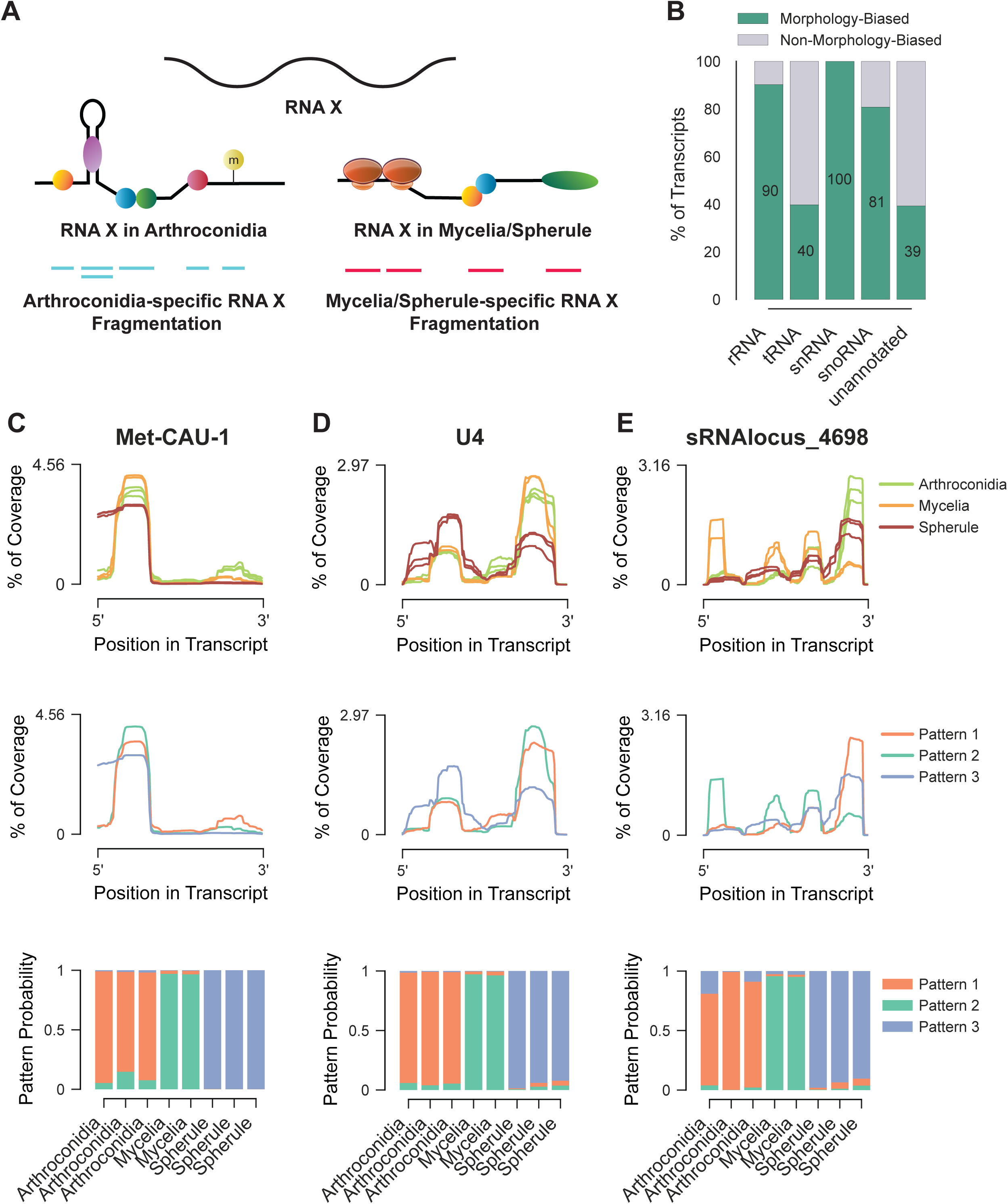
Analysis of RNA fragmentation patterns in different *Coccidioides* morphologies. A) Schematic representation of an RNA exhibiting differential fragmentation patterns in arthroconidia versus mycelia/spherules forms. B) Bar chart depicting the percentage of transcripts from different RNA classes, with and without significant morphology bias (padj < 0.05). C-E) Graphs showing the percentage of coverage across the transcript positions for three different RNA samples: Met-CAU-1 (C), U4 (D), and sRNALocus_4698 (E) in arthroconidia (orange), mycelia (green), and spherules (blue) forms, along with three modeled patterns of fragmentation (middle). Below each coverage graph, bar charts represent the calculated probability of observing each of the three fragmentation patterns across different morphological states.

Given the myriad of regulatory elements life cycle transitions between saprobic to parasitic which may alter the fragmentation of *Coccidioides* RNAs, we compared RiboMarker®-enhanced small RNA sequencing libraries of arthroconidia, mycelia, and spherule morphologies (Figure 2A). Briefly, ncRNAs identified using tRNAScanSE and RFAM, or unannotated sRNA-producing loci identified using ShortStack, were used for differential fragmentation profile analysis. Of these molecules we found that more often than not a transcript exhibited a fragmentation pattern significantly biased to different morphologies using our set of annotated and unannotated loci (Figure 4B) (34).

Three examples of these morphology-biased fragmentation profiles are shown in Figures 5C-5E. Figure 4C shows the RNA fragmentation profile from arthroconidia, mycelia, and spherules across tRNA-Methionine (Met)-CAU-1, a transfer RNA responsible for decoding of Methionine codons found in mRNA during protein translation and often involved in translation initiation (71). Transcript coverage (top panel) reveals differential fragmentation patterns near the 3’ end of the tRNA in both arthroconidia and mycelia morphologies, while absence of these fragments in spherule. Additionally, 5’ fragment coverage in the spherule is more robust starting at the 5’ terminal base compared to arthroconidia and mycelia, which show coverage beginning a few bases downstream. Using a machine-learning approach, we categorized these read pileups into three distinct patterns to capture these data (middle panel). Notably it appeared that these patterns were representative of the three distinct *Coccidioides* morphologies (top and middle). We sought to probabilistically determine if there was any association between the three morphologies and the three RNA fragmentation patterns by fitting each individual replicate to the representative model generated for these transcripts (top fit to middle). Pattern membership probabilities were generated for each sample (bottom panel) where we observed perfect correlation between *Coccidioides* morphology and the associated RNA fragmentation patterns for tRNA-Met-CAU-1. Similarly, we observed this phenomenon regarding the U4 snRNA is shown in Figure 4D. Here we saw an increased complexity of fragmentation compared to tRNA-Met-CAU-1, where the ratio of 5’ and 3’ derived fragments modulates relative to morphology type. In spherules, we observe a more even distribution of fragments derived from the 5’ and 3’ end of U4, while arthroconidia and mycelia show more 3’ bias to fragment generation (Figure 4D; top panel). Again, these differential fragmentation profiles allow us to delineate between replicates based on associated morphologies, suggesting these profiles uniquely represent their associated morphology (Figure 4D; bottom panel). Morphology-specific fragmentation patterns are not specific to annotated loci, as is shown in the expressed, unannotated RNA sRNAlocus_4698 (Figure 4E). Here, mycelia presents a unique fragmentation pattern with the generation of 5’ end and internal fragments as compared to arthroconidia and spherule, which display various amounts of more 3’ end and 3’ proximal fragments. Again, these patterns can be isolated and are found to show morphological specificity (Figure 4E; middle and bottom panels). Overall, while it cannot be said in any way that fragmentation of these highlighted RNAs (and other) elicit morphological changes in *Coccidioides*, their correlation with specific morphologies may be a tool in honing in on morphology-specific sRNAs and RNA fragments for use as RNA biomarkers downstream, especially those unique or highly abundant in the spherule, parasitic morphology.

**Figure 5.**
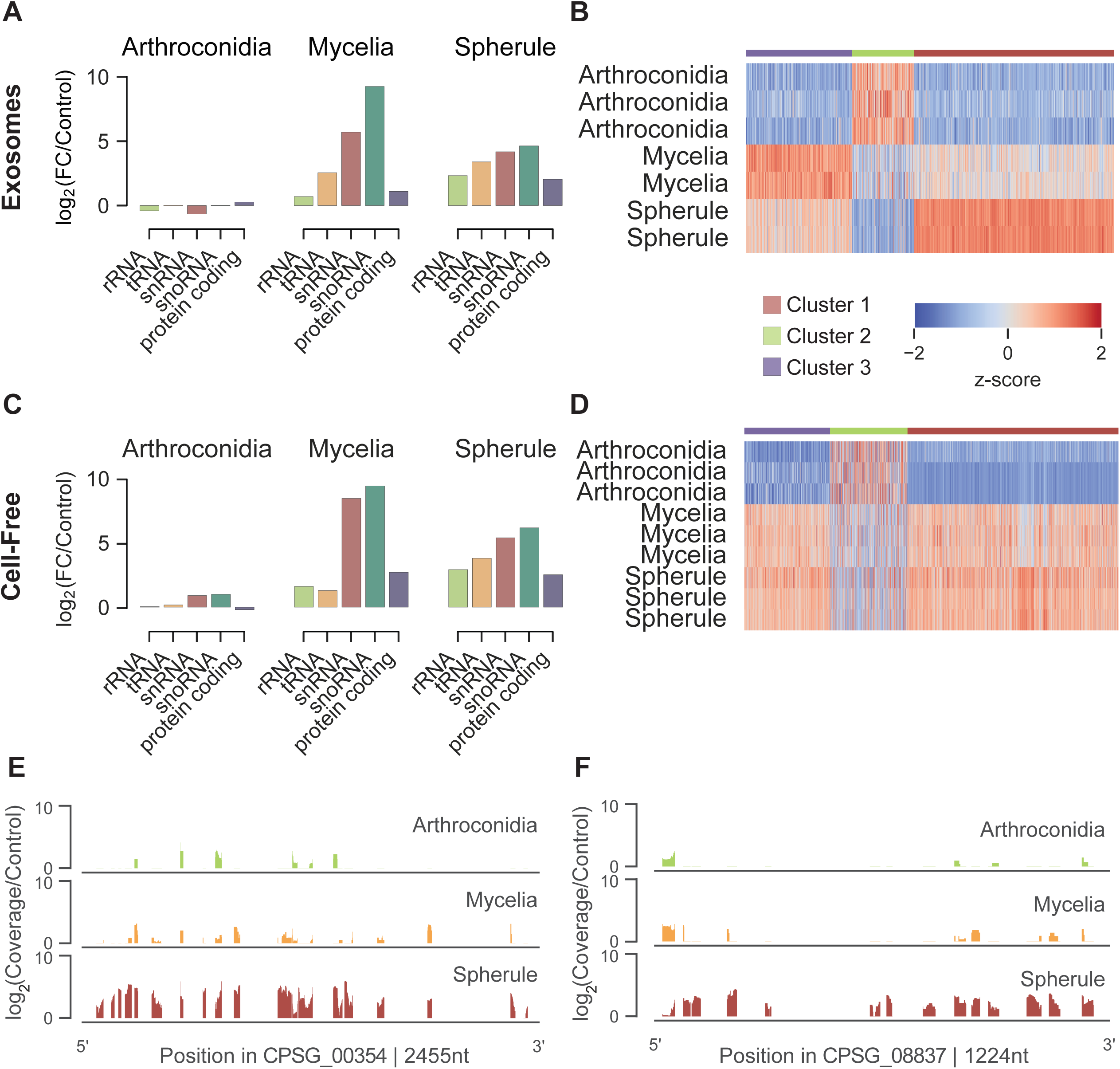
Extracellular RNA profiles vary across *C. posadasii* morphologies. **(A, C)** Bar plots showing the enrichment over control of small RNA reads mapping to annotated RNA-generating loci for exosomal **(A)** and cell-free **(C)** fractions of conditioned media across *Coccidioides* morphologies. Clustered heatmaps of small RNA sequencing data from RiboMarker® libraries derived from exosome-**(B)** and cell-free-**(D)** associated fractions. The exosome heatmap represents 1,239 RNAs, and the cell-free heatmap represents 2,651 RNAs, showing significant abundance differences across time points and spherule samples (padj < 0.01). **(E, F)** Read coverage plots for two predicted genes, CPSG_00354 **(E**; predicted Heat Shock protein 70) and CPSG_08837 **(F**; predicted Enolase), illustrating the mapped read coverage of extracellular RNAs from exosomal samples at different stages.

### Cell-free and exosomal small RNA pools contain putative effectors of cell-cell and pathogen-host communication

Upon infection, fungal pathogens can modulate the host immune response to generate a preferred microenvironment to promote pathogenesis (20). This is often achieved through the export of molecular effectors (e.g. proteins, RNAs, etc.) packaged in exosomes, a sub-population of extracellular vesicles with the ability to interact and fuse with lipid bilayers (72), facilitating cross-kingdom communication with host cells (26). Exported RNA effectors are known to, in part, target RNAi machinery in host cells to selectively silence target genes (28), which can lead to a dampened or neutralized host response, allowing the fungal pathogen to persist and propagate within the host (73). While these processes are well known in plant-associated fungal pathogens (74), little is known about the RNA content of the extracellular environment of *Coccidioides.* Therefore, we sought to leverage sequencing data from our extracellular fractions to determine what, if any, RNA exists, or may be actively exported, outside of the cell, and how this may change across *Coccidioides* morphologies.

To explore the potential of extracellular RNA related to *C. posadasii*, we performed RiboMarker® sequencing on exosome- and cell-free-associated RNAs isolated from fractions of filtered, conditioned media used in the growth and propagation of the arthroconidia, mycelia, and spherules samples. To control for background RNA not derived from the fungus, we prepared media-only (Control) samples for both exosomal and cell-free RNA fractions at all morphological stages (see *Methods*). An initial analysis of detected RNA expression across annotated sRNA gene types suggested large differences in the extracellular RNA pool from cell-free fractions of arthroconidia versus the mycelia and (Supplemental Figure 10; top scatter plot). As well as a larger presence of protein-coding derived RNAs in the exosomal spherule RNA pools relative to the mycelia (Supplemental Figure 10; bottom scatter plot), suggesting a large increase in the export of protein-coding RNA via exosomes to the extracellular space. To investigate the high-degree of similarity in the RNAs detected in either the cell-free or exosomal fractions (Supplemental Figure 10) we performed a transcript level analysis (Supplemental Figure 11) where we found the two fractions were highly correlated with one another (r>0.9), suggesting there are minimal discernable characteristics between the cell-free and exosomal RNA pools.

Relative to our media controls, the distribution of RNA types across our different time points varied significantly (Figure 5A/C). Among mycelia and spherules, we saw a >80-fold increase in snRNA-derived RNAs in the cell-free fraction and exosome fractions (Figure 5A/C), with U2 snRNA-derived RNAs seeing the highest mean enrichment relative to their respective controls. This marked extracellular increase may be indicative of enhanced intracellular splicing activity and could be associated with global transcriptomic reprogramming as the fungus prepares for further growth and/or differentiation (75). In the subsequent transition from the mycelia to spherules, we noted a substantial shift: the abundance of snRNA-derived reads dropped by half, while there was a notable increase in tRNA-derived reads (Figure 5C), which was also mirrored in the exosome fractions (Figure 5A). This increase in tRNA fragments might be indicative of stress response and adaptation mechanisms associated with environmental changes during formation of spherules or could reflect increased protein synthesis (70, 76).

Clustering of expression data from both cell-free and the exosome fractions suggested distinct RNA populations were significantly (padj<0.01) enriched relative to the *Coccidioides* morphology (Figure 5B/D). For the exosome samples, we saw a small population of RNAs enriched relative to the mycelia and spherules (Figure 5B; Cluster 1). Interestingly, a significant majority of extracellular small RNA and RNA fragments were found present in higher abundance in both the mycelia and spherule exosomal fractions, (compared to arthroconidia), with cluster 1 enriched in spherules and cluster 3 more abundant in the mycelia. (Figure 5B; Clusters 1/3; Supplemental File 5). The cell-free fraction highlighted similar patterns of RNA enrichment (Figure 5D; Supplemental File 6), with clusters 2 and 3 containing highly abundant molecules in both mycelia and spherules, whereas cluster 1 seems more highly enriched in arthroconidia; however, not as distinctly as the exosome fraction. Together these data suggest that the extracellular RNA content suggest that life cycle and metabolic states of *Coccidioides* may play a role in regulating RNA export *en masse*.

To gain more perspective on the biological function of the pool of extracellular RNAs, we focused on the exosome fraction, specifically those RNAs found specifically enriched in spherules (Cluster 1) (Figure 5B). In these data, we observed two intriguing examples of enriched small RNAs derived from protein-coding genes that have been previously shown in the literature to associate with extracellular vesicles (72). For example, small RNAs derived from CPSG_00354, encoding a predicted Hsp70-like protein, show an 8-fold increase in abundance in the spherule exosome fraction relative to the control and time course samples (Figure 5E). Interestingly, Hsp70 domain-containing protein motifs, found in protein chaperones and known virulence factors in pathogenic fungi (77, 78), have been shown to be the only universal factor found across extracellular vesicles tested from a panel of six different fungal pathogens (79), suggesting that heat-shock related proteins could play a universal role in fungal EV biogenesis and host interactions (77). Another example was the presence of small RNAs derived from a predicted enolase gene (Figure 5F) enriched in the spherule morphology. The glycolytic enzyme product of this protein-coding gene has been previously shown to be a major constituent of fungal extracellular vesicles (80) and has previously been used as a successful antigen in vaccination models in mice for *C. albicans* infections. It is important to note that we are only detecting associated mRNA fragments to these proteins, and proteins themselves can be loaded into EVs without mRNAs present. Whether protein translation occurs in extracellular vesicles is currently unknown; however, there is evidence that human exosomes transport mRNA fragments that may act as competing RNA to regulate gene expression in recipient cells (81). While more evidence would be required to confirm functions of mRNA-derived RNAs in *Coccidioides* extracellular vesicles, it offers a compelling hypothesis for these highlighted mRNA fragments and others we detected in this study.

Outside of mRNAs-derived small RNAs, we also found specific tRNA fragments enriched in our extracellular spherule samples, including Leucine tRNA and Proline tRNA fragments (Supplemental Figure 12). While their function outside of the cell is yet to be described, tRNA fragments like these have been shown to populate fungal extracellular vesicles (28), possibly playing roles in the regulation of host gene expression (82). More evidence is needed to determine the biological roles of these extracellular RNAs, outside of potential RNA biomarkers. Future directions could involve validating whether these extracellular RNAs, especially those enriched at certain life stages, play active roles in pathogenesis, signaling, or environmental adaptation in *Coccidioides*.

## DISCUSSION

We charted the small RNA transcriptome of three key morphological forms of *Coccidioides* encompassing both the saprobic and parasitic life cycles. We found that the application of our RiboMarker®sequencing across each morphological stage resulted in an increased incorporation of previously undetected small RNAs in *Coccidioides*, many of which derive from tRNAs, rRNAs, and RNAs derived from novel loci. In this same vein, our data has allowed us to validate, and expand, hundreds of predicted RNAs of previously described roles (e.g. miRNAs, snRNAs, etc.), as well as identify novel sRNA loci within the unannotated regions of the *C. posadasii* genome. These data also elucidate dynamic shifts in small RNA regulation across the *Coccidioides* life cycle, highlighting RNAs and RNA fragments that show enrichment in distinct morphologies, suggesting potential stage-specific functions or processing key to arthroconidia, mycelia, and spherules. Finally, our cell-free and exosome RNA sequencing results suggest a unique pool of RNA and RNA fragments make their way to the extracellular space. Together these data provide a first look at the morphology dependent changes in small RNA expression inside and outside the cell for *Coccidioides*.

Perhaps the most striking finding from this work was the number of unannotated RNA loci we discovered. ShortStack analysis uncovered nearly 500 RNA-expressing loci, many of which exhibited expression levels like that of annotated loci. Additionally, a subset appears to be morphology specific in their expression. While it is easy to accept that, like all life, *Coccidioides* also contains “dark matter” in its transcriptome, it is also reasonable that many of these are most likely unannotated non-coding RNAs, such as snoRNAs, snRNAs, or micro/microRNA-like RNAs. A MirDeep2 analysis of these datasets revealed no predicted miRNA-like genes (data not shown), which is concerning given this bioinformatics tool has been used previously to identify miRNAs and microRNA-like RNAs (milRNAs) from fungi. However, this does highlight the lack of bioinformatics tools for small RNA identification in fungi. While tools like *milRNAPredictor* (83) have been designed with milRNAs in mind, newer tools will be necessary to account for our present understanding of fungal small RNAs.

Beyond the innovative approaches to increasing inclusion of incompatible RNAs into the RealSeq-Biofluids small RNA library preparation through use of the RiboMarker® platform, we implemented a cutting-edge fragmentation analysis to ascertain if RNA fragmentation patterns of annotated and/or unannotated RNA species could be indicative of specific morphological states of *Coccidioides.* While this information may appear redundant in characterizing fungal samples of known life cycle states, this proof-of-concept may have far-reaching implications in the context of identifying viable RNA biomarkers for diagnostic purposes without the need for host sample sequencing or identification of unknown pathogens inflicting patients through a straightforward sequencing experiment. Additional experiments will need to be undertaken to determine if RNA fragmentation profiles across differing morphological states from a variety of species are as informative as they appear to be in *Coccidioides*.

Beyond morphological differences, our work aims to establish evidence for extracellular RNA profiles based on *Coccidioides* lifecycle stages. Fungal pathogens through evolution have established communication pathways with their hosts on the cellular level, through a dissemination of genetic and proteomic information. Extracellular vesicles play a role as key mediators of this cell-to-cell communication to regulate pathogenic processes (11). These processes are often to the advantage of the pathogen by suppressing the host immune response (12), establishing microenvironments suitable for pathogen growth and proliferation (13), and other yet undefined pathways. However, genetic information can also be disseminated through natural lysis of fungal cells and extracellular vesicles, releasing cellular RNA, DNA, and protein that can circulate in the host in a variety of biofluids. These RNAs, like host RNAs, are subject to degradation by nucleases and other agonists, but sometimes “stabilize” at a specific length, most likely due to their secondary structure, modifications, or RNA-binding protein-mediated protection. We hypothesize that through evolutionary processes many of the extracellular RNAs detected in this and future studies will not only be represented in this small RNA atlas for *Coccidioides* but will also represent the most robust RNAs or RNA fragments, suggesting biological functionality and reproducibility across infected hosts.

From a larger sequencing perspective, our data can serve as an example of issues that may need to be addressed when determining the best approach to sequencing to meet the needs of the experimental approach. While many commercial kits exist for all manner of RNA and small RNA sequencing, no one kit is sufficient for all manner of RNA types, largely due to incorrect end chemistry for adapter ligations or inhibitory modifications that block enzymatic processes of library preparation. Many protocols have been developed in the field of RNA research to either identify RNA species with specific end chemistries (84–86) or to increase inclusion of “hard to sequence” RNAs (42, 87–89). While results may vary depending on sample type, use of the RiboMarker® platform does reveal hidden layers of small RNA and RNA fragments, of which their importance is only starting to be realized. Especially in the context of RNA biomarkers, whether it be for infectious disease or cancer detection, the biological context of how or for what purpose RNA or RNA fragments are generated may become subordinate to simple “yes/no” detection results from patient samples for diagnostic and prognostic purposes.

In conclusion, our findings provide insights into the small RNA transcriptional programs underlying several distinct morphological forms of *Coccidioides* and highlight the differential expression and fragmentation of RNAs that accompany this fungal pathogen’s transition from saprobic to parasitic growth. On a broader scope, this study also provides proof of principle for the utility of the RealSeq RiboMarker® platform. We hope that these findings and the data generated will provide a resource and stepping-stone to combat the increasing disease incidence and expanding geographic range of Valley fever.

## AUTHORS CONTRIBUTIONS

J.H., A.M., R.C., T.W, C.N., S.K., AND S.B.S conceived and planned the experiments. J.H., R.C., AND T.W carried out the experiments. A.M. carried out bioinformatics analyses. J.H., R.C., and T.W, contributed to sample preparation. J.H., A.M., R.C., T.W, C.N., S.K., AND S.B.S contributed to the interpretation of the results. J.H and A.M. took the lead in writing the manuscript. All authors provided critical feedback and helped shape the research, analysis and manuscript.

## ACKNOWLEDGMENTS

C.J.N. acknowledges support from the National Institutes of Health (NIH) National Institute of General Medical Sciences (NIGMS) award R35GM124594, and from the Kamangar family in the form of an endowed chair to C.J.N. Fellowship support was provided to T.W. by the University of California, Merced and the University of California Office of the President.

## SUPPLEMENTAL FIGURE LEGENDS

**Supp Figure 1.**
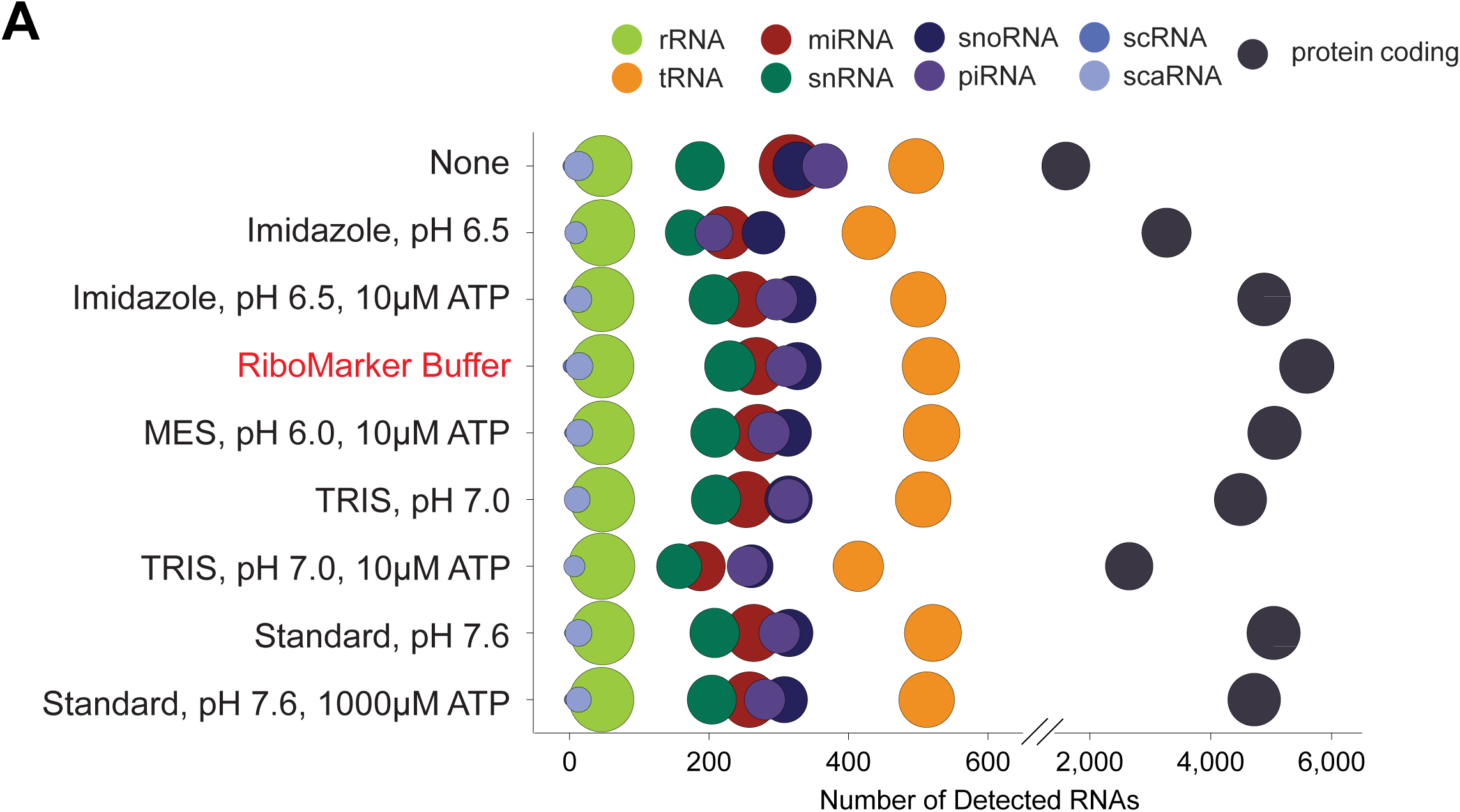
RiboMarker® Buffer captures the most diverse set of RNA molecules from the tested buffer conditions.

**Supp Figure 2.**
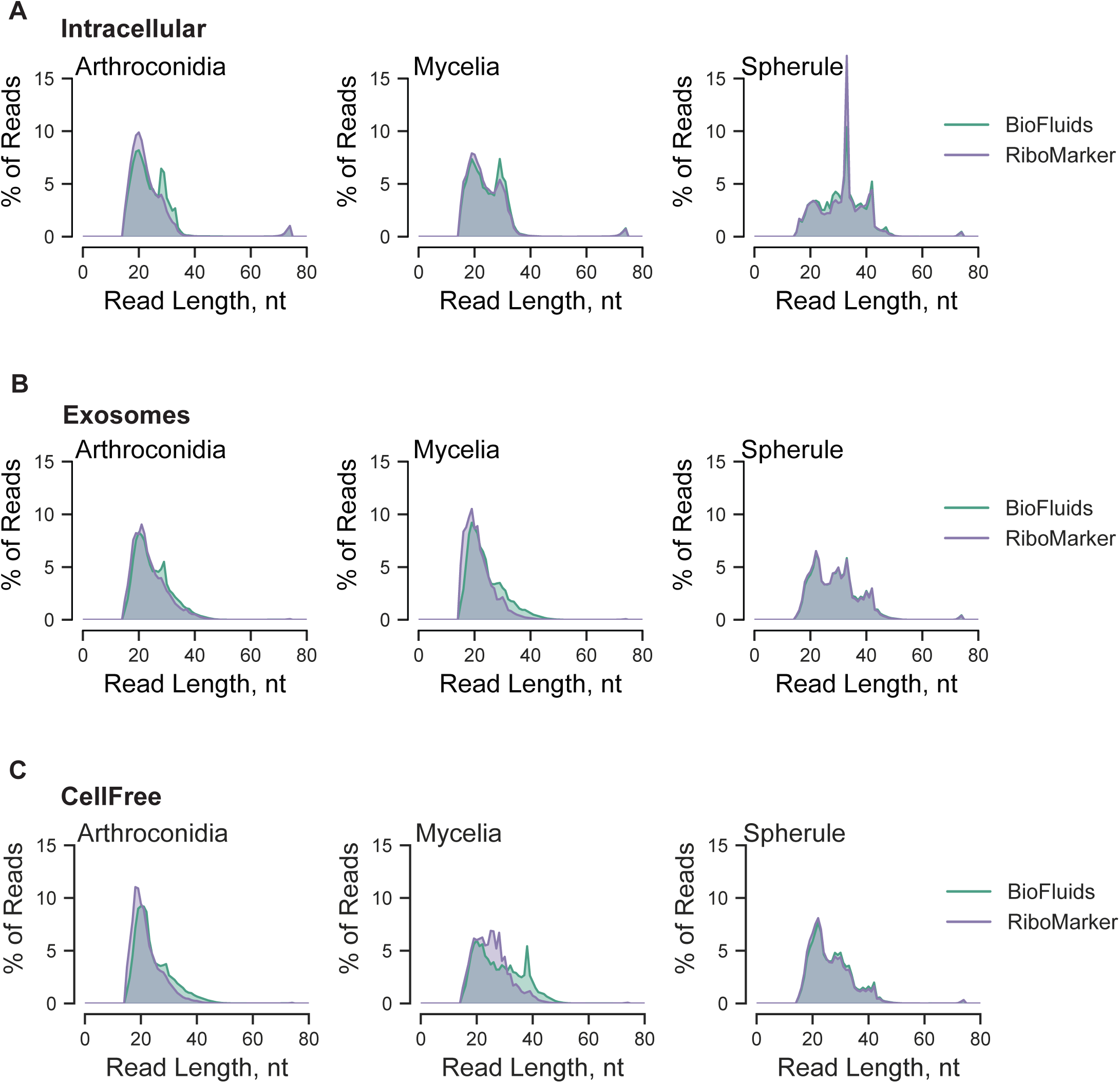
Read length distributions for reads mapping to *C. posadasii* for different morphologies and localizations.

**Supp Figure 3.**
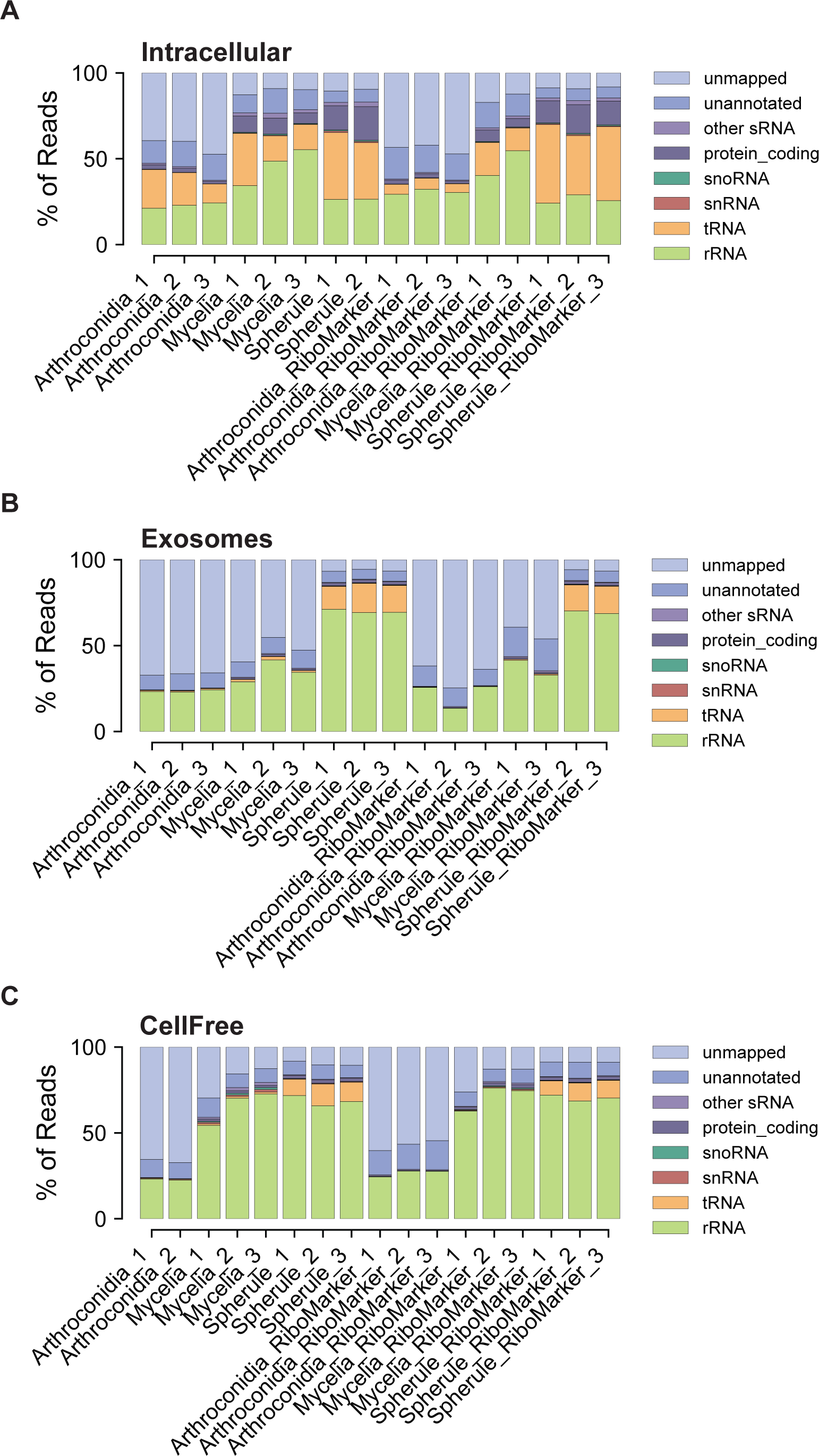
Breakdown of RNA transcript annotations for reads mapping to *C. posadasii* for different morphologies and localizations.

**Supp Figure 4.**
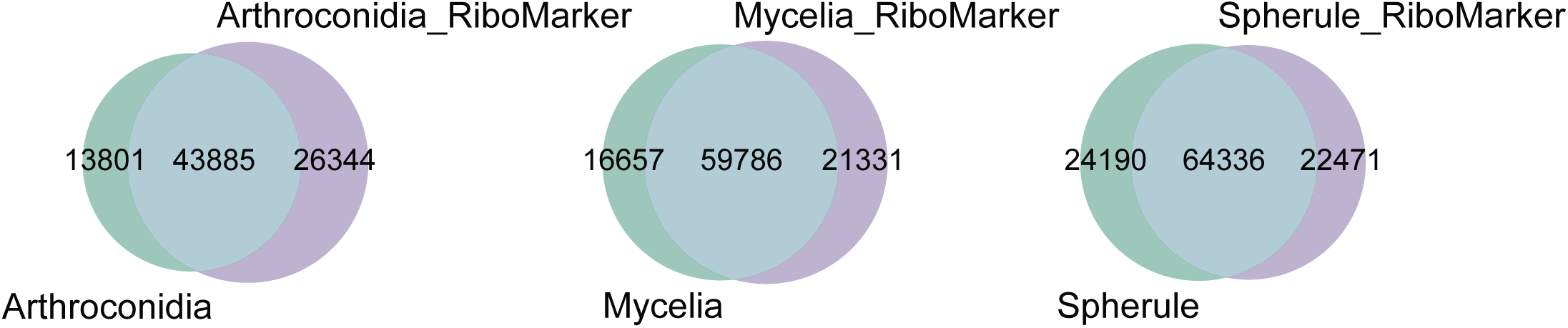
Differential sets of RNA transcripts are more readily incorporated into both Biofluids and RiboMarker® libraries.

**Supp Figure 5.**
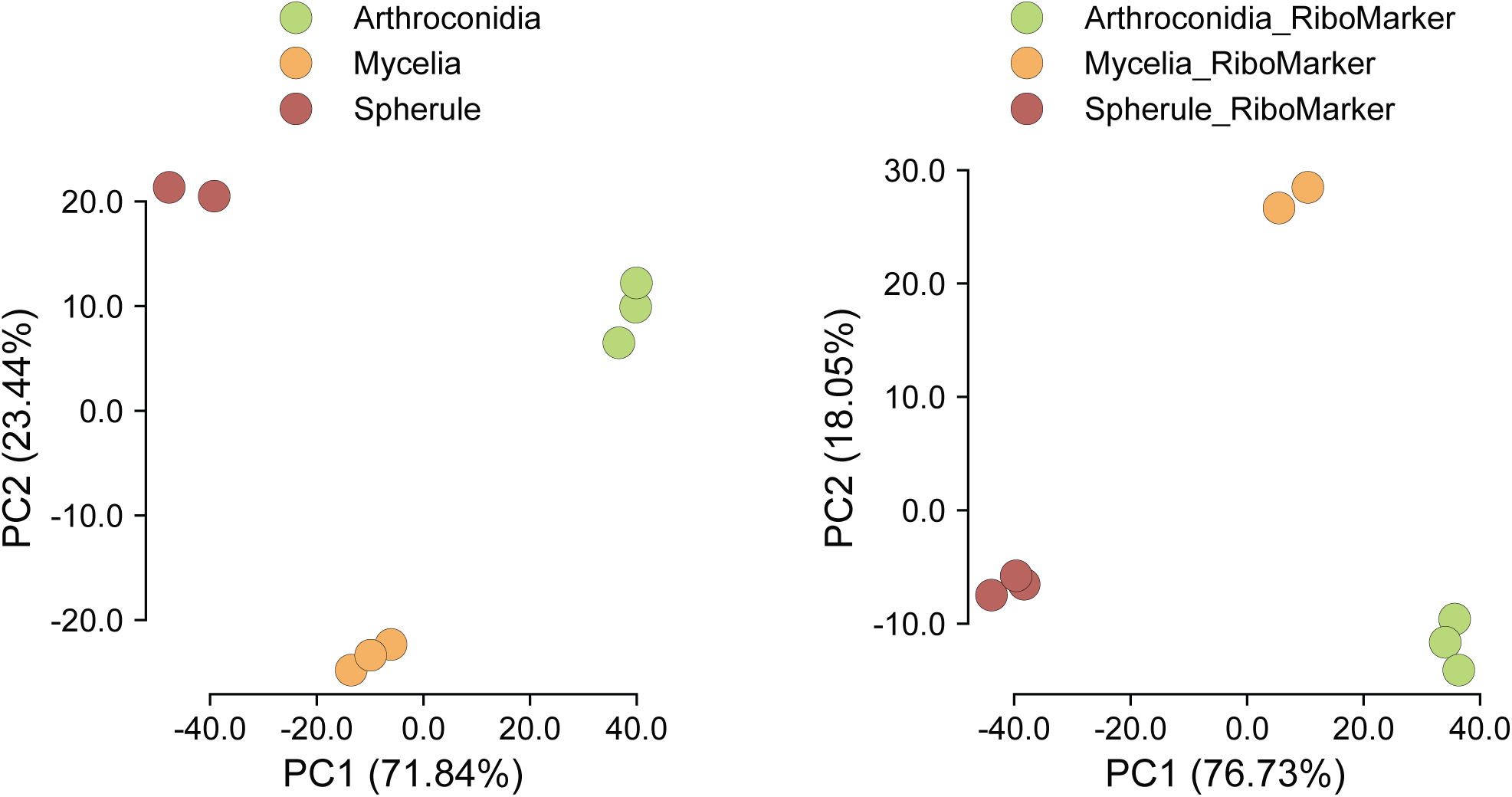
Principal component analysis of intracellular *C. posadasii* samples generated using Biofluids and RiboMarker®.

**Supp Figure 6.**
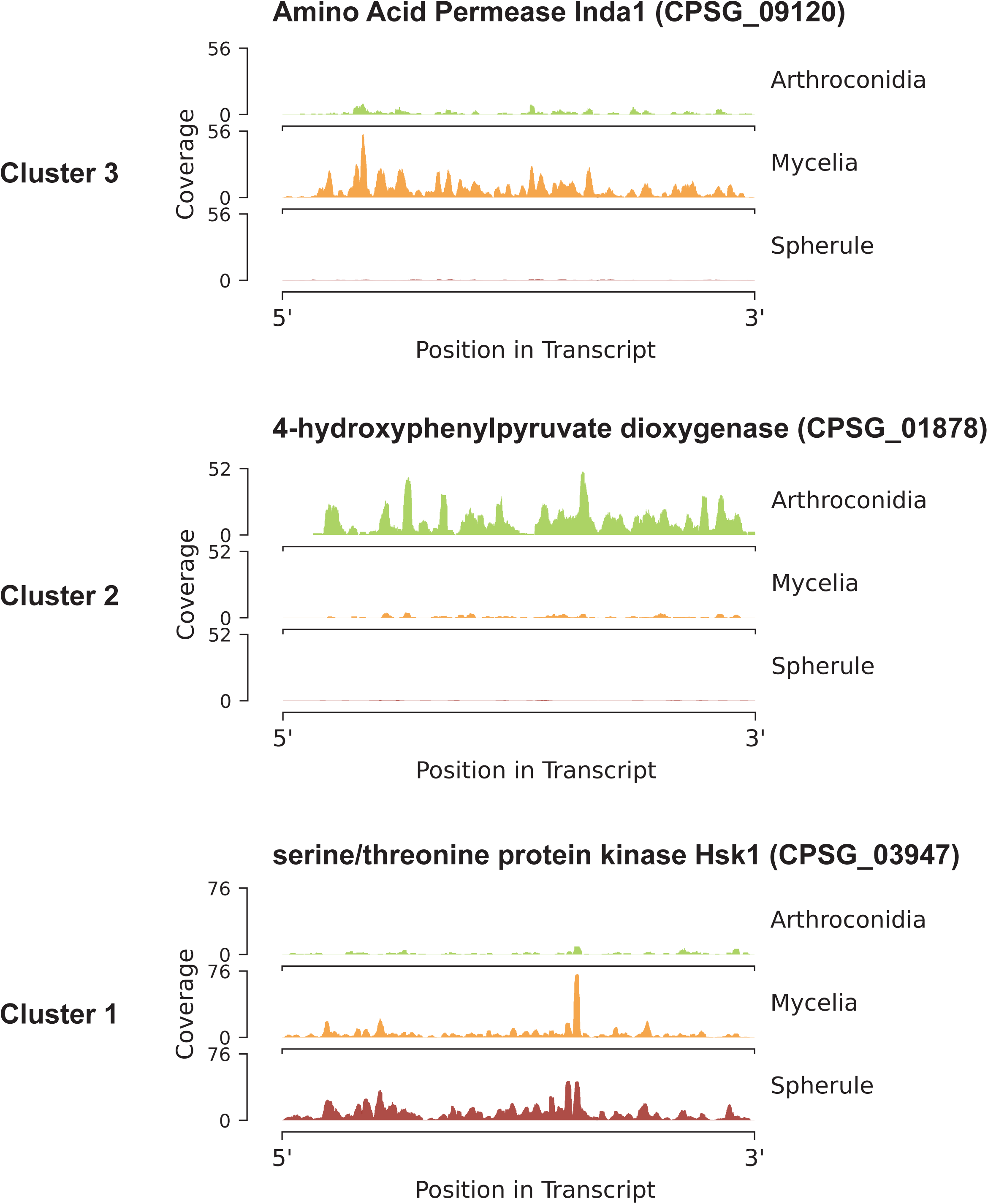
Read coverage across transcripts identified as differentially abundant across different morphological stages of *C. posadasii*.

**Supp Figure 7.**
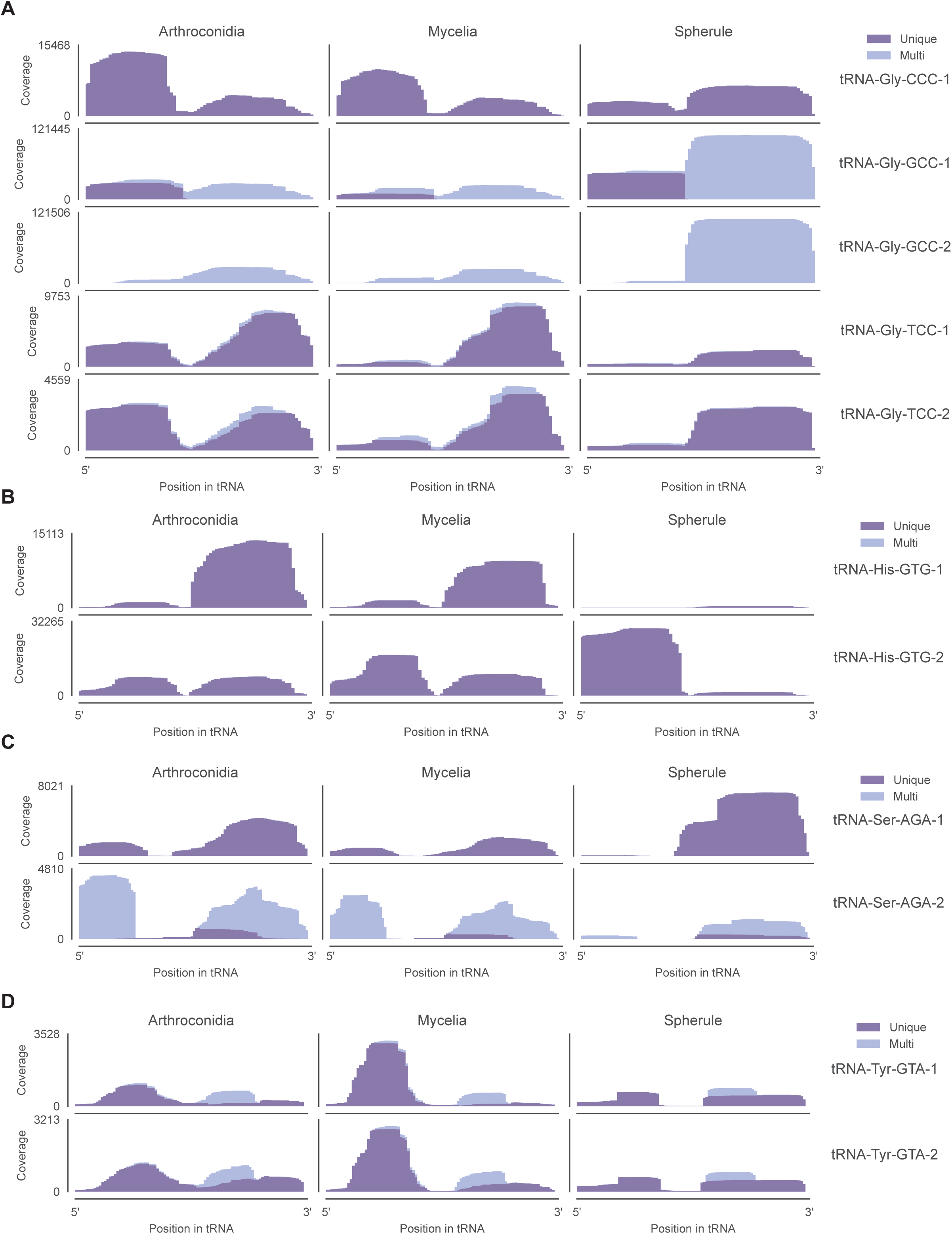
Read coverage across tRNA isodecoders that were identified as differentially abundant across different morphological stages of *C. posadasii.*Color is indicative of read mappability with dark purple being uniquely mapped to an individual tRNA isodecoder and light purple mapping to multiple different tRNA isodecoders.

**Supp Figure 8.**
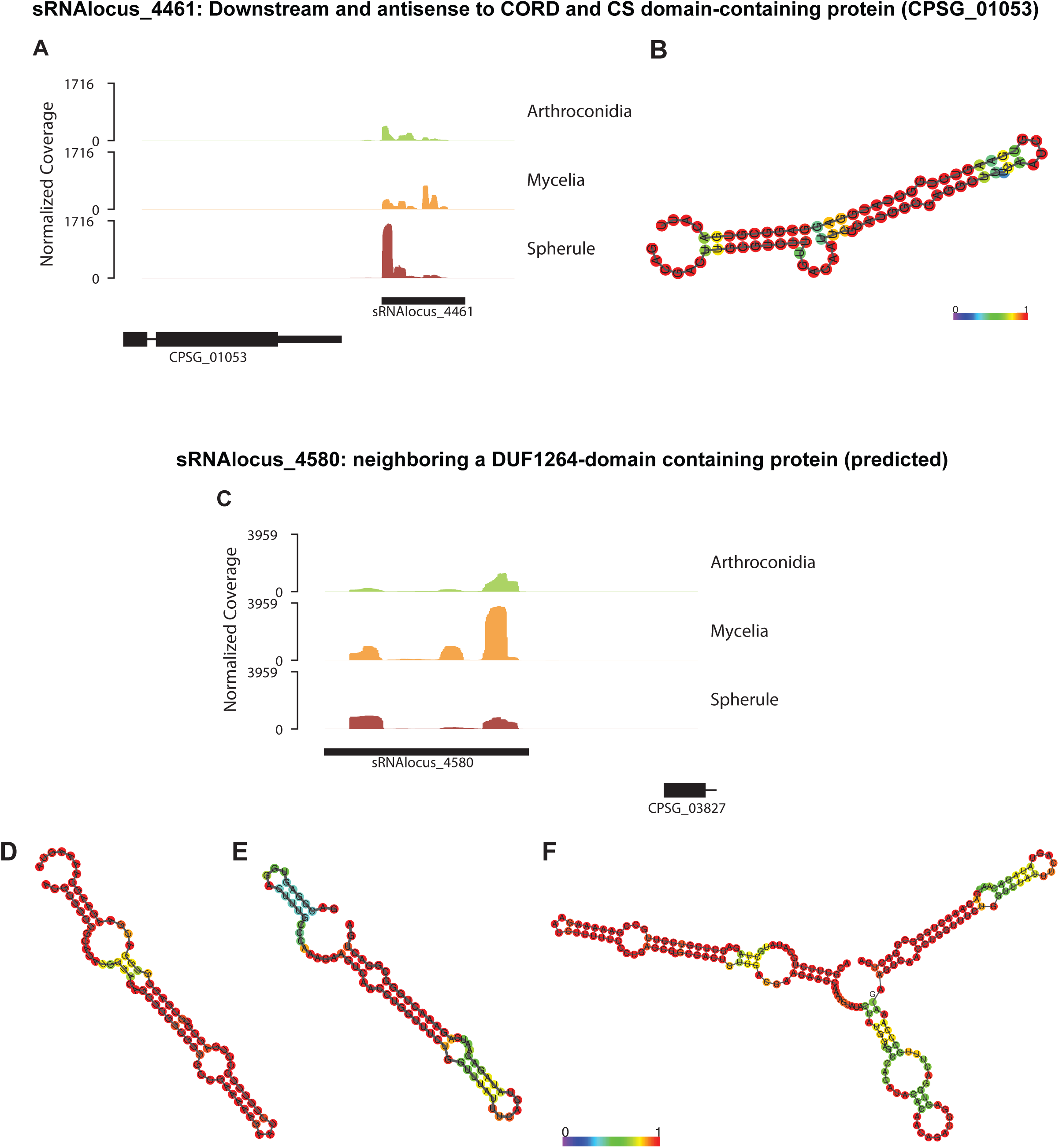
Read coverage of unannotated transcripts identified as differentially abundant across different morphological stages of *C. posadasii.* Predicted secondary structures based on minimum free energy folding are presented for each highlighted area (red boxes; **B,D,E**) and the entirety of sRNAlocus_4580 (**F).**

**Supp Figure 9.**
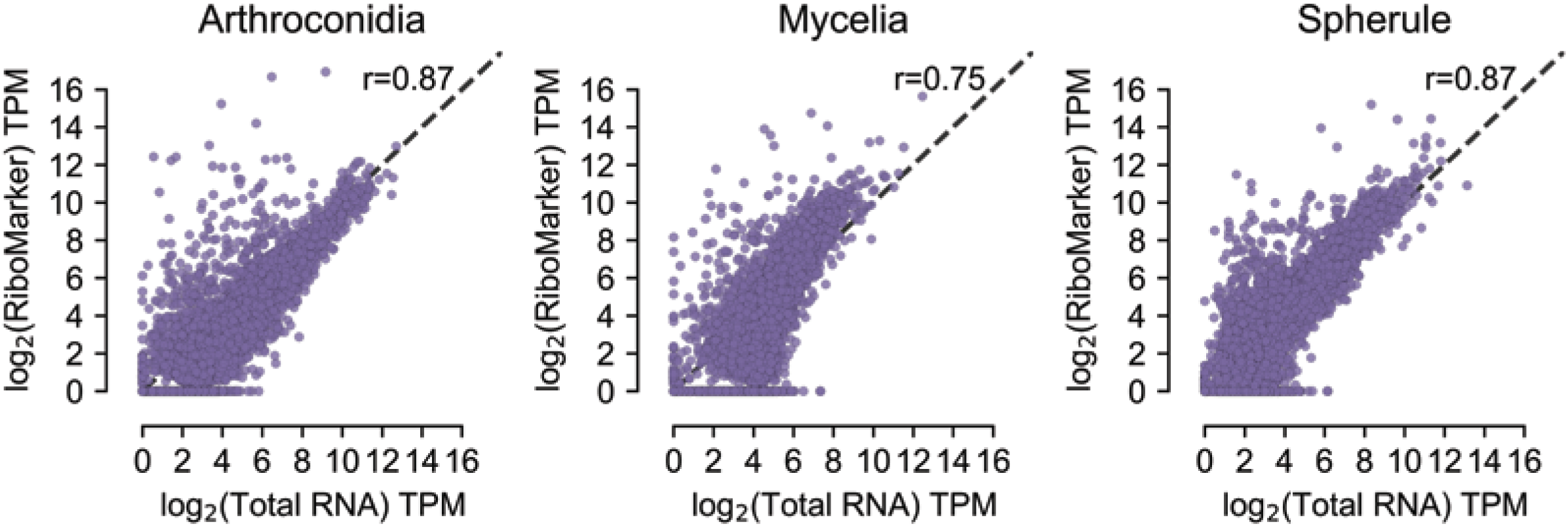
Scatter plot of log2(TPM) values for protein_coding mapped reads from RiboMarker (y-axis) versus those from the Zymo RiboFree Total RNA library preparation (x-axis). Pearson correlation coefficient values are included for each morphology.

**Supp Figure 10.**
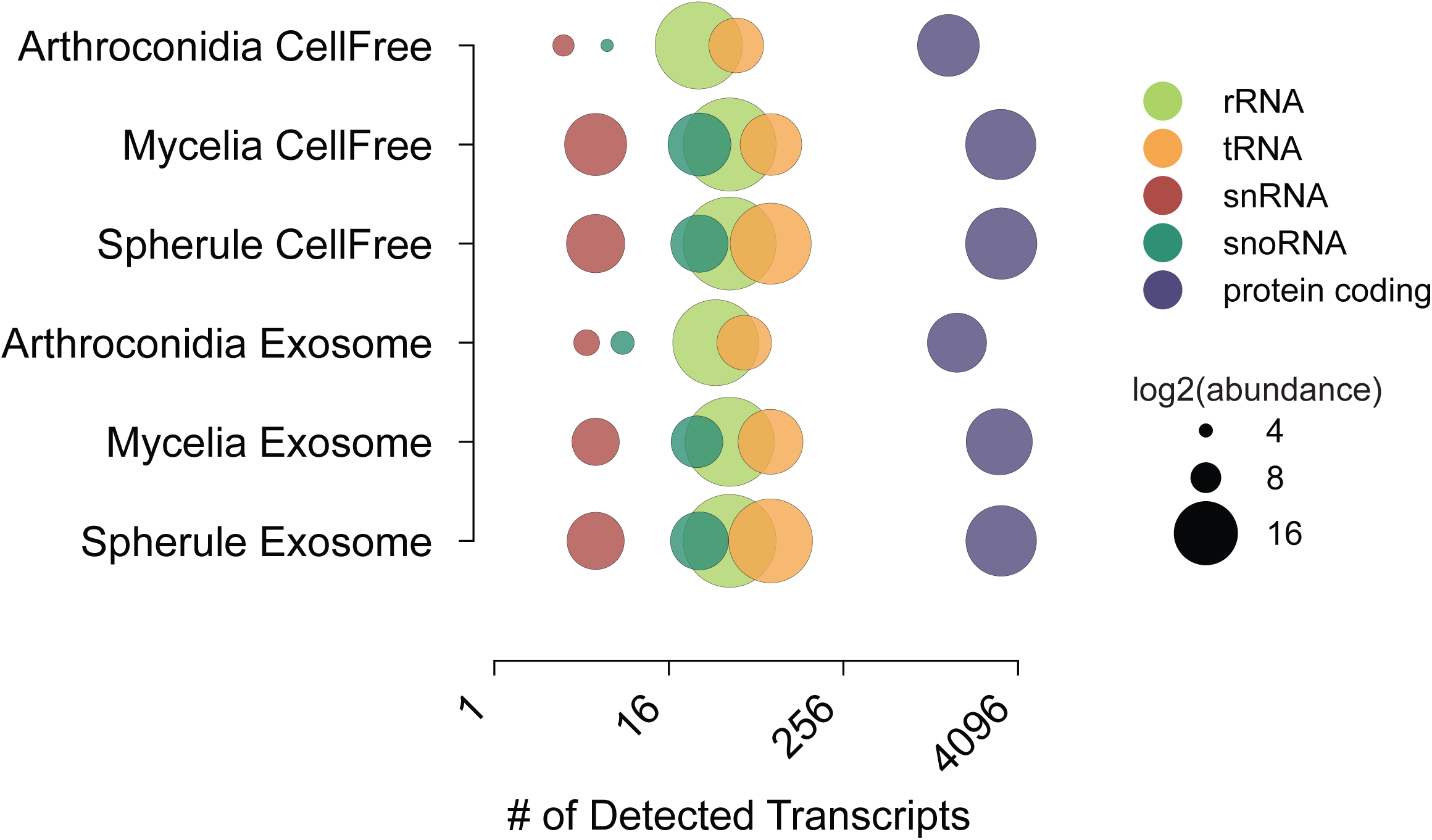
Dot plot representing the number of detected transcript annotations mapped to *C. posadasii* for cell-free **(top)** and exosome **(bottom)** samples using RiboMarker®, with dot size proportional to the log2(abundance) of each RNA class.

**Supp Figure 11.**
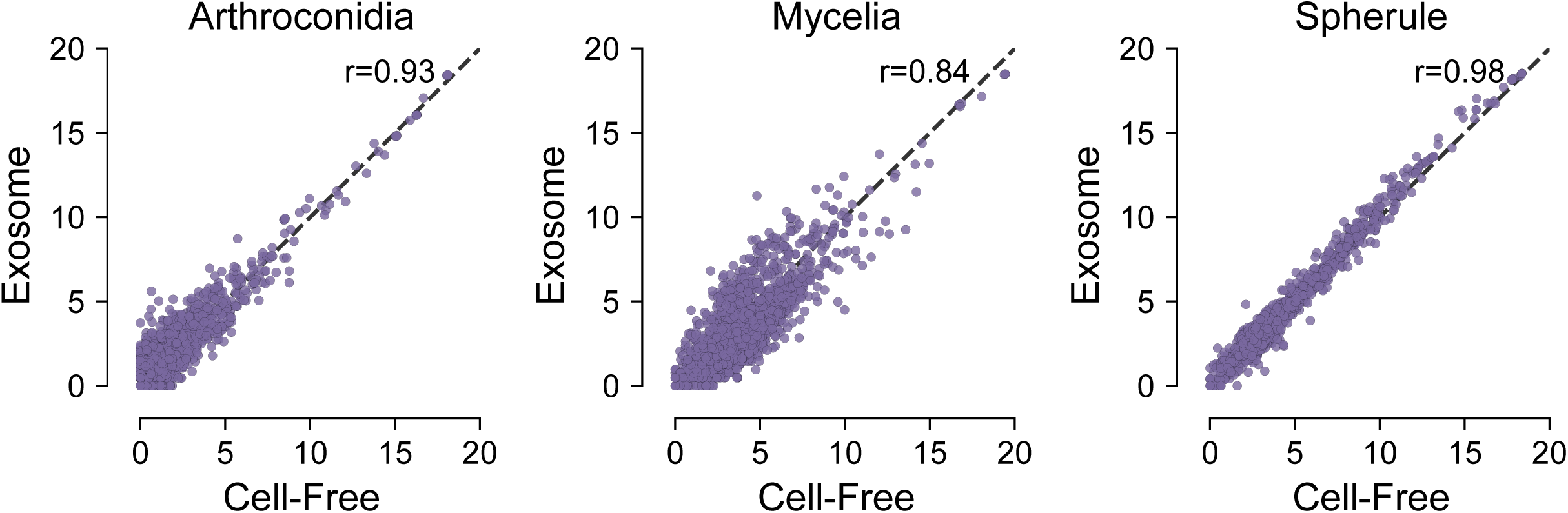
Comparative analysis of annotated transcript abundance between exosome and cell-free RNA preparations for arthroconidia,mycelia,, and spherule samples from *C. posadasii*. The Pearson correlation coefficient (r) for each comparison is shown at the top of each plot, indicating a strong correlation across all time points and sample types.

**Supp Figure 12.**
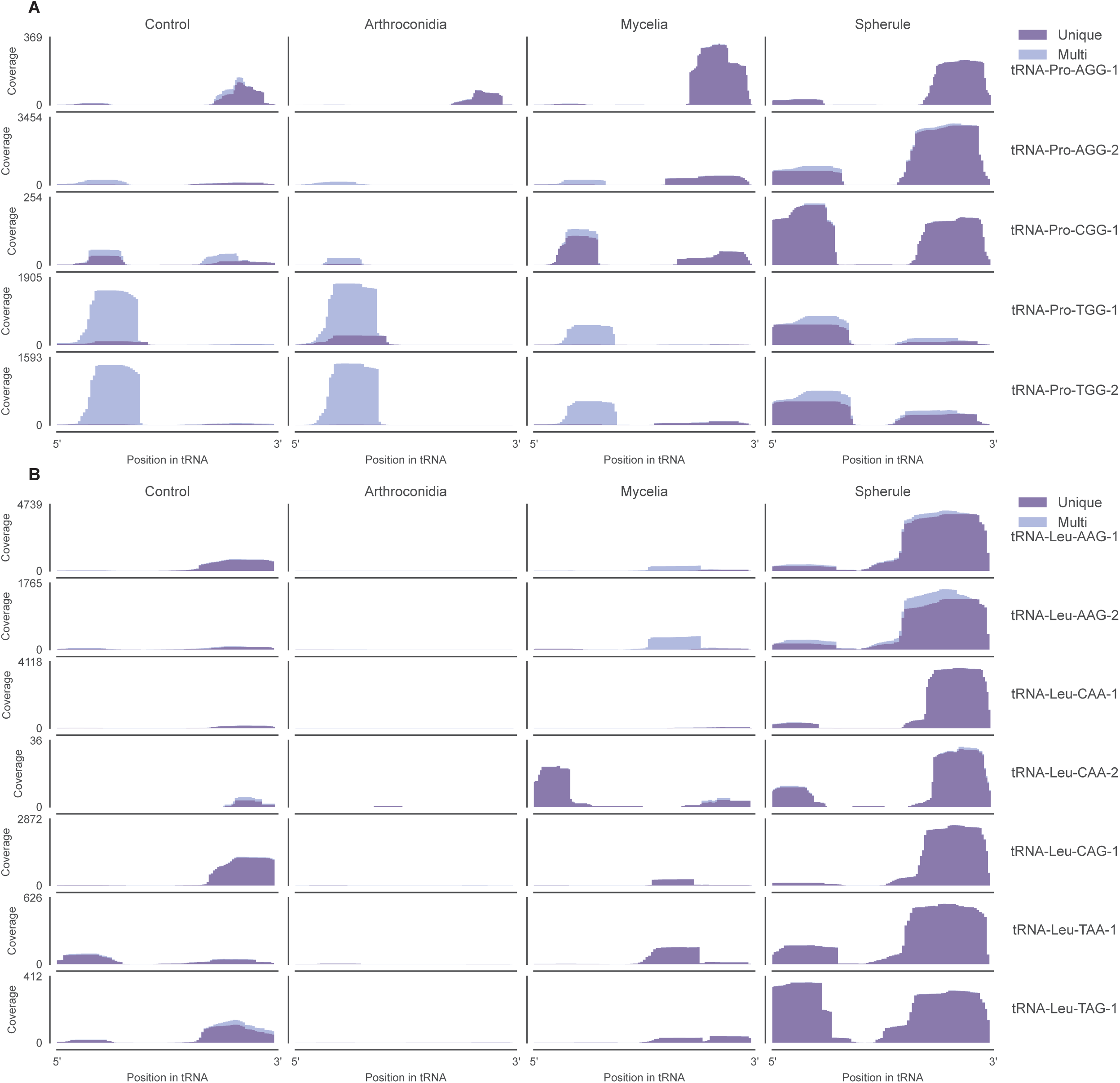
Read coverage across tRNA isodecoders that were identified as differentially abundant across exosome fractions of arthroconidia, mycelia, and spherule samples for *C. posadasii.* Color is indicative of read mappability with purple being uniquely mapped to an individual tRNA isodecoder and blue mapping to multiple different tRNA isodecoders.

## REFERENCES

1. Nguyen C, Barker BM, Hoover S, Nix DE, Ampel NM, Frelinger JA, et al. Recent advances in our understanding of the environmental, epidemiological, immunological, and clinical dimensions of coccidioidomycosis. Clin Microbiol Rev. 2013;26(3):505–25.

2. Facts and Stats about Valley Fever: U.S. Center for Disease Control and Prevention; [Available from: https://www.cdc.gov/valley-fever/php/statistics/index.html.

3. Galgiani JN, Ampel NM, Blair JE, Catanzaro A, Johnson RH, Stevens DA, et al. Coccidioidomycosis. Clin Infect Dis. 2005;41(9):1217–23.

4. Lockhart SR, Chowdhary A, Gold JAW. The rapid emergence of antifungal-resistant human-pathogenic fungi. Nat Rev Microbiol. 2023;21(12):818–32.

5. Odio CD, Marciano BE, Galgiani JN, Holland SM. Risk Factors for Disseminated Coccidioidomycosis, United States. Emerg Infect Dis. 2017;23(2):308–11.

6. Martinson ML, Lapham J. Prevalence of Immunosuppression Among US Adults. JAMA. 2024;331(10):880–2.

7. Williams SL, Chiller T. Update on the Epidemiology, Diagnosis, and Treatment of Coccidioidomycosis. J Fungi (Basel). 2022;8(7).

8. Gorris ME, Neumann JE, Kinney PL, Sheahan M, Sarofim MC. Economic Valuation of Coccidioidomycosis (Valley Fever) Projections in the United States in Response to Climate Change. Weather Clim Soc. 2021;13(1):107–23.

9. Organization WH. WHO fungal priority pathogens list to guide research, development and public health action: World Health Organization; 2022.

10. Carlin AF, Beyhan S, Pena JF, Stajich JE, Viriyakosol S, Fierer J, et al. Transcriptional Analysis of Coccidioides immitis Mycelia and Spherules by RNA Sequencing. J Fungi (Basel). 2021;7(5).

11. Narra HP, Shubitz LF, Mandel MA, Trinh HT, Griffin K, Buntzman AS, et al. A Coccidioides posadasii CPS1 Deletion Mutant Is Avirulent and Protects Mice from Lethal Infection. Infect Immun. 2016;84(10):3007–16.

12. Duttke SH, Beyhan S, Singh R, Neal S, Viriyakosol S, Fierer J, et al. Decoding Transcription Regulatory Mechanisms Associated with Coccidioides immitis Phase Transition Using Total RNA. mSystems. 2022;7(1):e0140421.

13. Mandel MA, Beyhan S, Voorhies M, Shubitz LF, Galgiani JN, Orbach MJ, et al. The WOPR family protein Ryp1 is a key regulator of gene expression, development, and virulence in the thermally dimorphic fungal pathogen Coccidioides posadasii. PLoS Pathog. 2022;18(4):e1009832.

14. Gou LT, Zhu Q, Liu MF. Small RNAs: An expanding world with therapeutic promises. Fundam Res. 2023;3(5):676–82.

15. Chen Q, Zhou T. Emerging functional principles of tRNA-derived small RNAs and other regulatory small RNAs. J Biol Chem. 2023;299(10):105225.

16. Lambert M, Benmoussa A, Provost P. Small Non-Coding RNAs Derived From Eukaryotic Ribosomal RNA. Noncoding RNA. 2019;5(1).

17. Zinder JC, Lima CD. Targeting RNA for processing or destruction by the eukaryotic RNA exosome and its cofactors. Genes Dev. 2017;31(2):88–100.

18. Qutob D, Chapman BP, Gijzen M. Transgenerational gene silencing causes gain of virulence in a plant pathogen. Nat Commun. 2013;4:1349.

19. Jin Y, Zhao JH, Zhao P, Zhang T, Wang S, Guo HS. A fungal milRNA mediates epigenetic repression of a virulence gene in Verticillium dahliae. Philos Trans R Soc Lond B Biol Sci. 2019;374(1767):20180309.

20. Lai Y, Jiang B, Hou F, Huang X, Ling B, Lu H, et al. The emerging role of extracellular vesicles in fungi: a double-edged sword. Front Microbiol. 2023;14:1216895.

21. Hua C, Zhao JH, Guo HS. Trans-Kingdom RNA Silencing in Plant-Fungal Pathogen Interactions. Mol Plant. 2018;11(2):235–44.

22. Islam W, Islam SU, Qasim M, Wang L. Host-Pathogen interactions modulated by small RNAs. RNA Biol. 2017;14(7):891–904.

23. Katiyar-Agarwal S, Jin H. Role of small RNAs in host-microbe interactions. Annu Rev Phytopathol. 2010;48:225–46.

24. He B, Wang H, Liu G, Chen A, Calvo A, Cai Q, et al. Fungal small RNAs ride in extracellular vesicles to enter plant cells through clathrin-mediated endocytosis. Nat Commun. 2023;14(1):4383.

25. Cui C, Wang Y, Liu J, Zhao J, Sun P, Wang S. A fungal pathogen deploys a small silencing RNA that attenuates mosquito immunity and facilitates infection. Nat Commun. 2019;10(1):4298.

26. Cheng AP, Lederer B, Oberkofler L, Huang L, Johnson NR, Platten F, et al. A fungal RNA-dependent RNA polymerase is a novel player in plant infection and cross-kingdom RNA interference. PLoS Pathog. 2023;19(12):e1011885.

27. Mahanty B, Mishra R, Joshi RK. Cross-kingdom small RNA communication between plants and fungal phytopathogens-recent updates and prospects for future agriculture. RNA Biol. 2023;20(1):109–19.

28. Bitencourt TA, Pessoni AM, Oliveira BTM, Alves LR, Almeida F. The RNA Content of Fungal Extracellular Vesicles: At the “Cutting-Edge” of Pathophysiology Regulation. Cells. 2022;11(14).

29. Mead HL, Van Dyke MCC, Barker BM. Proper Care and Feeding of Coccidioides: A Laboratorian’s Guide to Cultivating the Dimorphic Stages of C. immitis and C. posadasii. Curr Protoc Microbiol. 2020;58(1):e113.

30. Converse JL, Besemer AR. Nutrition of the Parasitic Phase of Coccidioides Immitis in a Chemically Defined Liquid Medium. J Bacteriol. 1959;78(2):231–9.

31. Martin M. Cutadapt removes adapter sequences from high-throughput sequencing reads. 2011. 2011;17(1):3.

32. Chan PP, Lin BY, Mak AJ, Lowe TM. tRNAscan-SE 2.0: improved detection and functional classification of transfer RNA genes. Nucleic Acids Res. 2021;49(16):9077–96.

33. Axtell MJ. ShortStack: comprehensive annotation and quantification of small RNA genes. RNA. 2013;19(6):740–51.

34. Antonio Cuevas MF, Ricardo Fraiman. An anova test for functional data. Computational Statistics & Data Analysis. 2004;47(1):111–22.

35. Lai H, Feng N, Zhai Q. Discovery of the major 15-30 nt mammalian small RNAs, their biogenesis and function. Nat Commun. 2023;14(1):5796.

36. Novogrodsky A, Hurwitz J. The enzymatic phosphorylation of ribonucleic acid and deoxyribonucleic acid. I. Phosphorylation at 5’-hydroxyl termini. J Biol Chem. 1966;241(12):2923–32.

37. Cameron V, Uhlenbeck OC. 3’-Phosphatase activity in T4 polynucleotide kinase. Biochemistry. 1977;16(23):5120–6.

38. Cameron V, Soltis D, Uhlenbeck OC. Polynucleotide kinase from a T4 mutant which lacks the 3’ phosphatase activity. Nucleic Acids Res. 1978;5(3):825–33.

39. Kollath DR, Miller KJ, Barker BM. The mysterious desert dwellers: Coccidioides immitis and Coccidioides posadasii, causative fungal agents of coccidioidomycosis. Virulence. 2019;10(1):222–33.

40. Xue J, Chen X, Selby D, Hung CY, Yu JJ, Cole GT. A genetically engineered live attenuated vaccine of Coccidioides posadasii protects BALB/c mice against coccidioidomycosis. Infect Immun. 2009;77(8):3196–208.

41. Fu M, Gu J, Wang M, Zhang J, Chen Y, Jiang P, et al. Emerging roles of tRNA-derived fragments in cancer. Mol Cancer. 2023;22(1):30.

42. Hernandez R, Shi J, Liu J, Li X, Wu J, Zhao L, et al. PANDORA-Seq unveils the hidden small noncoding RNA landscape in atherosclerosis of LDL receptor-deficient mice. J Lipid Res. 2023;64(4):100352.

43. Klotz SA, Drutz DJ, Huppert M, Sun SH, DeMarsh PL. The critical role of CO2 in the morphogenesis of Coccidioides immitis in cell-free subcutaneous chambers. J Infect Dis. 1984;150(1):127–34.

44. Cappellazzo G, Lanfranco L, Fitz M, Wipf D, Bonfante P. Characterization of an amino acid permease from the endomycorrhizal fungus Glomus mosseae. Plant Physiol. 2008;147(1):429–37.

45. Wipf D, Benjdia M, Tegeder M, Frommer WB. Characterization of a general amino acid permease from Hebeloma cylindrosporum. FEBS Lett. 2002;528(1-3):119–24.

46. Shimmoto M, Matsumoto S, Odagiri Y, Noguchi E, Russell P, Masai H. Interactions between Swi1-Swi3, Mrc1 and S phase kinase, Hsk1 may regulate cellular responses to stalled replication forks in fission yeast. Genes Cells. 2009;14(6):669–82.

47. Ward RJ, Angert ER. DNA replication during endospore development in Metabacterium polyspora. Mol Microbiol. 2008;67(6):1360–70.

48. Wang Y, Wang Z, Jibril SM, Wei M, Pu X, Yang C, et al. Transcriptome and Quasi-Targeted Metabolome Analyze Overexpression of 4-Hydroxyphenylpyruvate Dioxygenase Alleviates Fungal Toxicity of 9-Phenanthrol in Magnaporthe oryzae. Int J Mol Sci. 2022;23(13).

49. Qian H, Sun L, Wu M, Zhao W, Liu M, Liang S, et al. The COPII subunit MoSec24B is involved in development, pathogenicity and autophagy in the rice blast fungus. Front Plant Sci. 2022;13:1074107.

50. Viriyakosol S, Singhania A, Fierer J, Goldberg J, Kirkland TN, Woelk CH. Gene expression in human fungal pathogen Coccidioides immitis changes as arthroconidia differentiate into spherules and mature. BMC Microbiol. 2013;13:121.

51. Torrent M, Chalancon G, de Groot NS, Wuster A, Madan Babu M. Cells alter their tRNA abundance to selectively regulate protein synthesis during stress conditions. Sci Signal. 2018;11(546).

52. Liu B, Cao J, Wang X, Guo C, Liu Y, Wang T. Deciphering the tRNA-derived small RNAs: origin, development, and future. Cell Death Dis. 2021;13(1):24.

53. Kapranov P, St Laurent G. Dark Matter RNA: Existence, Function, and Controversy. Front Genet. 2012;3:60.

54. Argaman L, Hershberg R, Vogel J, Bejerano G, Wagner EG, Margalit H, et al. Novel small RNA-encoding genes in the intergenic regions of Escherichia coli. Curr Biol. 2001;11(12):941–50.

55. Gervais NC, Shapiro RS. Discovering the hidden function in fungal genomes. Nat Commun. 2024;15(1):8219.

56. Dogra N, Chen TY, Gonzalez-Kozlova E, Miceli R, Cordon-Cardo C, Tewari AK, et al. Extracellular vesicles carry transcriptional ‘dark matter’ revealing tissue-specific information. J Extracell Vesicles. 2024;13(8):e12481.

57. Zhang M, Kadota Y, Prodromou C, Shirasu K, Pearl LH. Structural basis for assembly of Hsp90-Sgt1-CHORD protein complexes: implications for chaperoning of NLR innate immunity receptors. Mol Cell. 2010;39(2):269–81.

58. The RC. RNAcentral: a hub of information for non-coding RNA sequences. Nucleic Acids Res. 2019;47(D1):D1250–D1.

59. Sayers EW, Beck J, Bolton EE, Brister JR, Chan J, Comeau DC, et al. Database resources of the National Center for Biotechnology Information. Nucleic Acids Res. 2024;52(D1):D33–D43.

60. Lorenz R, Bernhart SH, Honer Zu Siederdissen C, Tafer H, Flamm C, Stadler PF, et al. ViennaRNA Package 2.0. Algorithms Mol Biol. 2011;6:26.

61. Maggio-Hall LA, Lyne P, Wolff JA, Keller NP. A single acyl-CoA dehydrogenase is required for catabolism of isoleucine, valine and short-chain fatty acids in Aspergillus nidulans. Fungal Genet Biol. 2008;45(3):180–9.

62. Tello-Ruiz MK, Jaiswal P, Ware D. Gramene: A Resource for Comparative Analysis of Plants Genomes and Pathways. Methods Mol Biol. 2022;2443:101–31.

63. Lee HC, Li L, Gu W, Xue Z, Crosthwaite SK, Pertsemlidis A, et al. Diverse pathways generate microRNA-like RNAs and Dicer-independent small interfering RNAs in fungi. Mol Cell. 2010;38(6):803–14.

64. Sonawane AR, Platig J, Fagny M, Chen CY, Paulson JN, Lopes-Ramos CM, et al. Understanding Tissue-Specific Gene Regulation. Cell Rep. 2017;21(4):1077–88.

65. Ando D, Rashad S, Begley TJ, Endo H, Aoki M, Dedon PC, et al. tRNA modifications inform tissue specific mRNA translation and codon optimization. bioRxiv. 2024:2023.10.24.563884.

66. Gilbert WV, Nachtergaele S. mRNA Regulation by RNA Modifications. Annu Rev Biochem. 2023;92:175–98.

67. Jonkhout N, Tran J, Smith MA, Schonrock N, Mattick JS, Novoa EM. The RNA modification landscape in human disease. RNA. 2017;23(12):1754–69.

68. Ule J, Jensen K, Mele A, Darnell RB. CLIP: a method for identifying protein-RNA interaction sites in living cells. Methods. 2005;37(4):376–86.

69. Underwood JG, Uzilov AV, Katzman S, Onodera CS, Mainzer JE, Mathews DH, et al. FragSeq: transcriptome-wide RNA structure probing using high-throughput sequencing. Nat Methods. 2010;7(12):995–1001.

70. Li G, Manning AC, Bagi A, Yang X, Gokulnath P, Spanos M, et al. Distinct Stress-Dependent Signatures of Cellular and Extracellular tRNA-Derived Small RNAs. Adv Sci (Weinh). 2022;9(17):e2200829.

71. Cigan AM, Feng L, Donahue TF. tRNAi(met) functions in directing the scanning ribosome to the start site of translation. Science. 1988;242(4875):93–7.

72. Liebana-Jordan M, Brotons B, Falcon-Perez JM, Gonzalez E. Extracellular Vesicles in the Fungi Kingdom. Int J Mol Sci. 2021;22(13).

73. Liu J, Hu X. Fungal extracellular vesicle-mediated regulation: from virulence factor to clinical application. Front Microbiol. 2023;14:1205477.

74. Zhou Q, Ma K, Hu H, Xing X, Huang X, Gao H. Extracellular vesicles: Their functions in plant-pathogen interactions. Mol Plant Pathol. 2022;23(6):760–71.

75. Muzafar S, Sharma RD, Chauhan N, Prasad R. Intron distribution and emerging role of alternative splicing in fungi. FEMS Microbiol Lett. 2021;368(19).

76. Nawrot B, Sochacka E, Duchler M. tRNA structural and functional changes induced by oxidative stress. Cell Mol Life Sci. 2011;68(24):4023–32.

77. Silveira CP, Piffer AC, Kmetzsch L, Fonseca FL, Soares DA, Staats CC, et al. The heat shock protein (Hsp) 70 of Cryptococcus neoformans is associated with the fungal cell surface and influences the interaction between yeast and host cells. Fungal Genet Biol. 2013;60:53–63.

78. Horianopoulos LC, Kronstad JW. Chaperone Networks in Fungal Pathogens of Humans. J Fungi (Basel). 2021;7(3).

79. Parreira V, Santos LGC, Rodrigues ML, Passetti F. ExVe: The knowledge base of orthologous proteins identified in fungal extracellular vesicles. Comput Struct Biotechnol J. 2021;19:2286–96.

80. Nimrichter L, de Souza MM, Del Poeta M, Nosanchuk JD, Joffe L, Tavares Pde M, et al. Extracellular Vesicle-Associated Transitory Cell Wall Components and Their Impact on the Interaction of Fungi with Host Cells. Front Microbiol. 2016;7:1034.

81. Batagov AO, Kurochkin IV. Exosomes secreted by human cells transport largely mRNA fragments that are enriched in the 3’-untranslated regions. Biol Direct. 2013;8:12.

82. Peres da Silva R, Puccia R, Rodrigues ML, Oliveira DL, Joffe LS, Cesar GV, et al. Extracellular vesicle-mediated export of fungal RNA. Sci Rep. 2015;5:7763.

83. Yao Y, Zhang H, Deng H. milRNApredictor: Genome-free prediction of fungi milRNAs by incorporating k-mer scheme and distance-dependent pair potential. Genomics. 2020;112(3):2233–40.

84. Honda S, Morichika K, Kirino Y. Selective amplification and sequencing of cyclic phosphate-containing RNAs by the cP-RNA-seq method. Nat Protoc. 2016;11(3):476–89.

85. Peach SE, York K, Hesselberth JR. Global analysis of RNA cleavage by 5’-hydroxyl RNA sequencing. Nucleic Acids Res. 2015;43(17):e108.

86. Kugelberg U, Natt D, Skog S, Kutter C, Ost A. 5 XP sRNA-seq: efficient identification of transcripts with and without 5 phosphorylation reveals evolutionary conserved small RNA. RNA Biol. 2021;18(11):1588–99.

87. Cozen AE, Quartley E, Holmes AD, Hrabeta-Robinson E, Phizicky EM, Lowe TM. ARM-seq: AlkB-facilitated RNA methylation sequencing reveals a complex landscape of modified tRNA fragments. Nat Methods. 2015;12(9):879–84.

88. Zheng G, Qin Y, Clark WC, Dai Q, Yi C, He C, et al. Efficient and quantitative high-throughput tRNA sequencing. Nat Methods. 2015;12(9):835–7.

89. Upton HE, Ferguson L, Temoche-Diaz MM, Liu XM, Pimentel SC, Ingolia NT, et al. Low-bias ncRNA libraries using ordered two-template relay: Serial template jumping by a modified retroelement reverse transcriptase. Proc Natl Acad Sci U S A. 2021;118(42).

